# Synaptic learning rules for sequence learning

**DOI:** 10.1101/2020.04.13.039826

**Authors:** Eric T. Reifenstein, Richard Kempter

## Abstract

Remembering the temporal order of a sequence of events is a task easily performed by humans in everyday life, but the underlying neuronal mechanisms are unclear. This problem is particularly intriguing as human behavior often proceeds on a time scale of seconds, which is in stark contrast to the much faster millisecond time-scale of neuronal processing in our brains. One long-held hypothesis in sequence learning suggests that a particular temporal fine-structure of neuronal activity — termed “phase precession” — enables the compression of slow behavioral sequences down to the fast time scale of the induction of synaptic plasticity. Using mathematical analysis and computer simulations, we find that — for short enough synaptic learning windows — phase precession can improve temporal-order learning tremendously and that the asymmetric part of the synaptic learning window is essential for temporal-order learning. To test these predictions, we suggest experiments that selectively alter phase precession or the learning window and evaluate memory of temporal order.

## Introduction

It is a pivotal quality for animals to be able to store and recall the order of events (“temporal-order learning”, 1–3) but there is only little work on the neural mechanisms generating asymmetric memory associations across behavioral time intervals (4). Putative mechanisms need to bridge the gap between the faster time scale of the induction of synaptic plasticity (typically milliseconds) and the slower time scale of behavioral events (seconds or slower). The slower time scale of behavioral events is mirrored, for example, in the time course of firing rates of hippocampal place cells (5), which signal when an animal visits certain locations (“place fields”) in the environment. The faster time scale is given by the temporal properties of the induction of synaptic plasticity (6, 7) — and spike-timing dependent plasticity (STDP) is a common form of synaptic plasticity that depends on the millisecond timing and temporal order of presynaptic and postsynaptic spiking. For STDP, the so-called “learning window” describes the temporal intervals at which presynaptic and postsynaptic activity induce synaptic plasticity. Such precisely timed neural activity can be generated by phase precession, which is the successive across-cycle shift of spike phases from late to early with respect to a background oscillation (Fig. 1). As an animal explores an environment, phase precession can be observed in the activity of hippocampal place cells with respect to the theta oscillation (8, 9). Phase precession is highly significant in single trials (10, 11) and occurs even in first traversals of a place field in a novel environment (12). Interestingly, phase precession allows for a temporal compression of a sequence of behavioral events from the time scale of seconds down to milliseconds (Fig. 1; 12–14), which matches the widths of generic STDP learning windows (15–18). This putative advantage of phase precession for temporal-order learning, however, has not yet been quantified. To assess the benefit of phase precession for temporal-order learning, we determine the synaptic weight change between pairs of cells whose activity represents two events of a sequence. Using both analytical methods and numerical simulations, we find that phase precession can dramatically facilitate temporal-order learning by increasing the synaptic weight change and the signal-to-noise ratio by up to an order of magnitude. We thus provide a mechanistic description of associative chaining models (19) and extend these models to explain how to store serial order.

**Figure 1:**
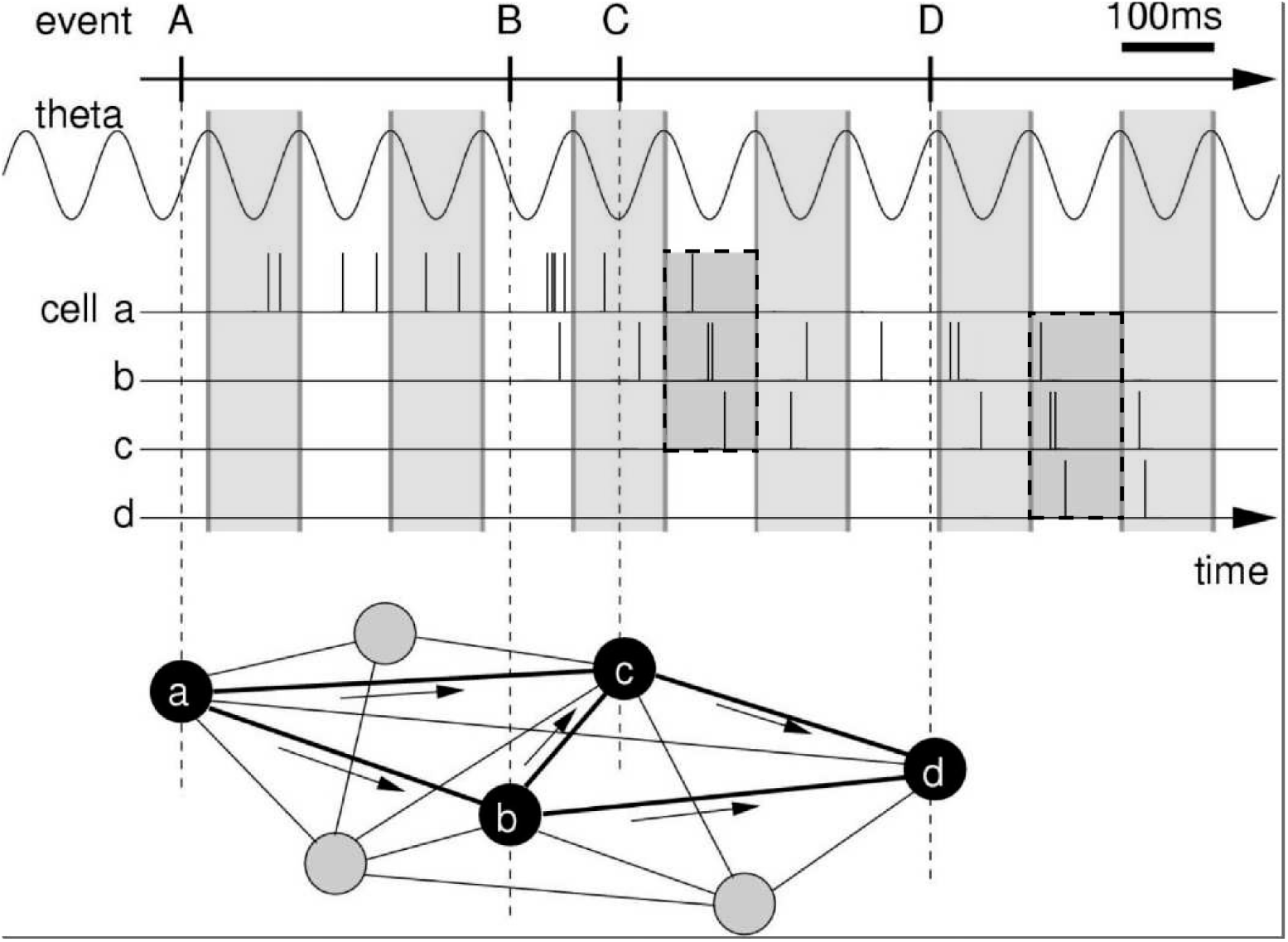
Rationale for temporal-order learning via phase precession. **Top**: Behavioral events (A to D) happen on a time scale of seconds. Middle: These events are represented by different cells (a-d), which fire a burst of stochastic action potentials in response to the onset of their respective event. We assume that each cell shows phase precession with respect to the LFP’s theta oscillation (every second cycle is marked by a grey box). When the activities of multiple cells overlap, the sequence of behavioral events is compressed in time to within one theta cycle (two examples highlighted in the dashed, shaded boxes). **Bottom**: This faster time scale can be picked up by STDP and strengthen the connections between the cells of the sequence.

## Results

To address the question of how behavioral sequences could be encoded in the brain, we study the change of synapses between neurons that represent events in a sequence. We assume that the temporal order of two events is encoded in the asymmetry of the efficacies of synapses that connect neurons representing the two events (Fig. 1). After the successful encoding of a sequence, a neuron that was activated earlier in the sequence has a strengthened connection to a neuron that was activated later in the sequence; whereas the connection in the reverse direction may be unchanged or is even weakened. As a result, when the first event is encountered and/or the first neuron is activated, the neuron representing the second event is activated. Consequently, the behavioral sequence could be replayed (as illustrated by simulations e.g., in 14, 20–26) and the memory of the temporal order of events is recalled (27, 28). We note, however, that in what follows we do not simulate such a replay of sequences, which would depend also on a vast number of parameters that define the network; instead, we rather focus on the underlying change in connectivity, which is the very basis of replay, and draw connections to “replay” in the *Discussion*.

Let us now illustrate key features of the encoding of the temporal order of sequences. To do so, we consider the weight change induced by the activity of two sequentially activated cells *i* and *j* that represent two behavioral events (dashed lines in Fig. 2A). Classical Hebbian learning (29), where weight changes Δ*w*_*ij*_ depend on the product of the firing rates *f*_*i*_ and *f*_*j*_, is not suited for temporal-order learning because the weight change is independent of the order of cells:

**Figure 2:**
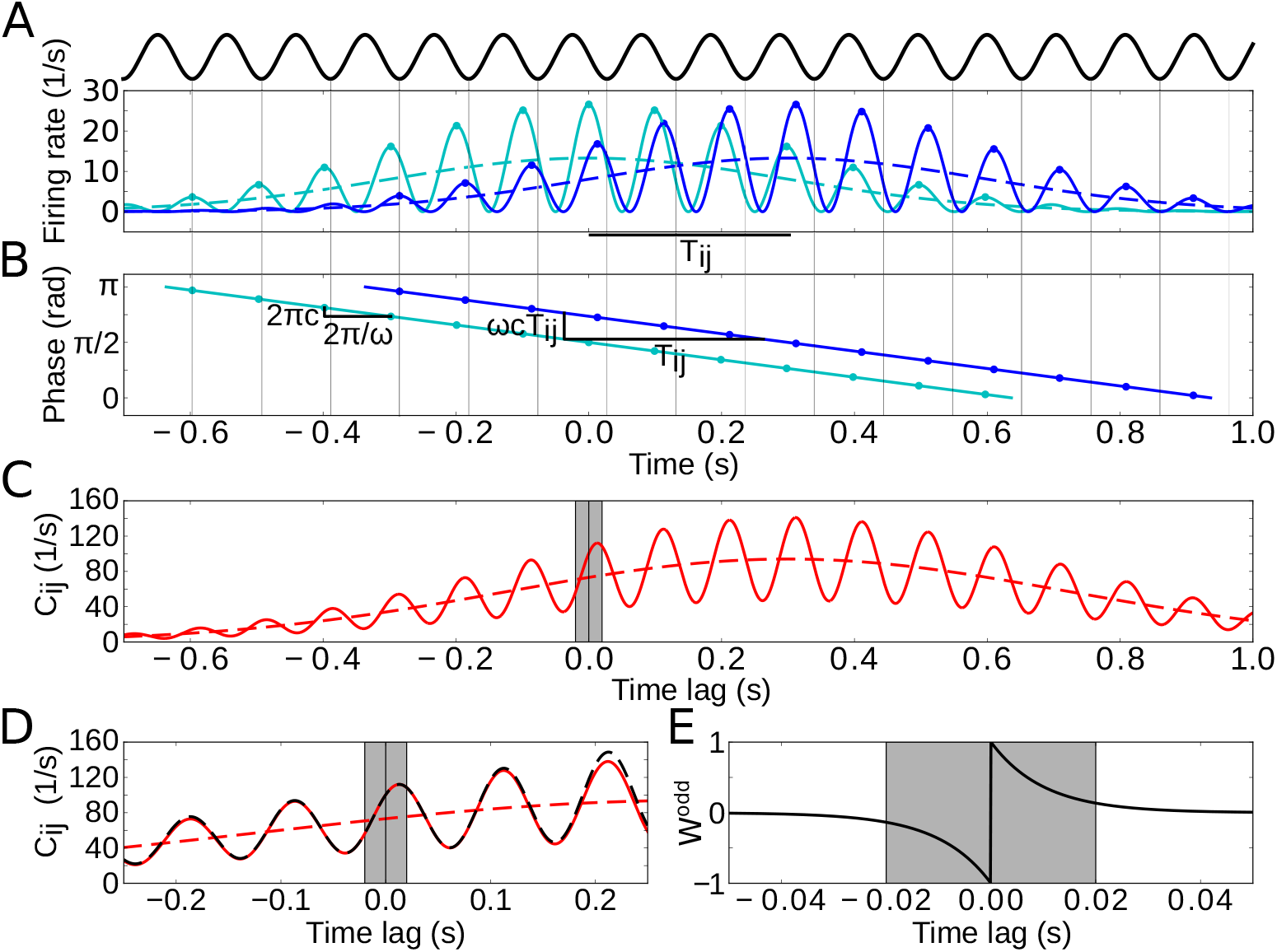
Model of two sequentially activated phase-precessing cells. **(A)** Oscillatory firing-rate profiles for two cells (solid blue and cyan lines). The black curve depicts the population theta oscillation. For easier comparison of the two different frequencies, the population activity’s troughs are continued by thin gray lines, and the peaks of the cell-intrinsic theta oscillation are marked by dots. Dashed lines depict the underlying Gaussian firing fields without theta modulation. **(B)** Phase precession of the two cells (same colors as in A). The compression factor *c* describes the phase shift per theta cycle for an individual cell (2*πc*). For the temporal separation *T*_*ij*_ of the firing fields and the theta frequency *ω*, the phase difference between the cells is *ωcT*_*ij*_. The dots depict the times of the maxima in (A). **(C)** Resulting cross-correlation for the two firing rates from (A). The solid red curve shows the full cross-correlation. The dashed line depicts the cross-correlation without theta-modulation. The grey region indicates small (< 20 ms) time lags. **(D)** Same as in (C), but zoomed in. Note that the first peak of the theta modulation is at a positive non-zero time lag, reflecting phase precession. The dashed black curve shows the approximation of the cross-correlation for the analytical treatment (Materials and Methods, Equation 17). **(E)** Synaptic learning window. The grey region indicates the region in which the learning window is large, and this region is also indicated in (C) and (D). Positive time lags correspond to postsynaptic activity following presynaptic activity. Parameters for all plots: *T*_*ij*_ = 0.3 s, *ω* = 2 *π ·* 10 Hz, *σ* = 0.3 s, *τ* = 10 ms, *µ* = 1, *c* = 0.042, *A* = 10.

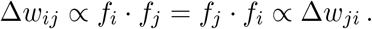

Therefore, a classical Hebbian weight change is symmetric, i.e., Δ*w*_*ij*_ − Δ*w*_*ji*_ = 0. This result can be generalized to learning rules that are based on the product of two arbitrary functions of the firing rates. We note that, although not suited for temporal-order learning, Hebbian rules are able to achieve more general “sequence learning”, where an association between sequence elements is created — independent of the order of events. To become sensitive to temporal order, we use spike-timing dependent plasticity (STDP; 6, 7). For STDP, average weight changes depend on the cross-correlation function of the firing rates (example in Fig. 2C,D),

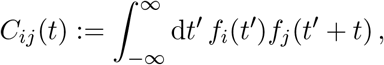

which is anti-symmetric: *C*_*ij*_(*t*) = *C*_*ji*_(−*t*). Assuming additive STDP, i.e., weight changes resulting from pairs of pre- and postsynaptic action potentials are added, the average synaptic weight change Δ*w*_*ij*_ between the two cells in a sequence can then be calculated explicitly (30):

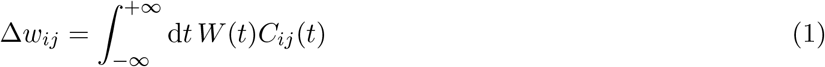

where *W* is the STDP learning window (example in Fig. 2E). We aim solve equation 1 for given firing rates *f*_*i*_ and *f*_*j*_. To do so, we assume that the synaptic weight *w*_*ij*_ is generally small and thus only has a weak impact on the cross-correlation of the cells during encoding, that is, for the “encoding” of a sequence the cross-correlation function is dominated by feedforward input whereas the recurrent inputs are neglected.

Next, let us show that the symmetry of *W* is essential for temporal-order learning. Any learning window *W* can be split up into an even part *W* ^even^, with *W* ^even^(*t*) = *W* ^even^(−*t*), and an odd part *W* ^odd^, with *W* ^odd^(*t*) = −*W* ^odd^(−*t*), such that *W* = *W* ^even^ + *W* ^odd^. For even learning windows, one can derive from Equation 1 and the anti-symmetry of *C*_*ij*_ that weight changes are symmetric, i.e. Δ*w*_*ij*_ = Δ*w*_*ji*_; therefore only the odd part *W* ^odd^ of *W* is useful for learning temporal order.

To further explore requirements for encoding the temporal order of a sequence of events, we restrict our analysis to odd learning windows. We then can relate the weight change Δ*w*_*ij*_ to the essential features of *C*_*ij*_(*t*). To do so, we integrate Equation 1 by parts (with *W* replaced by *W* ^odd^),

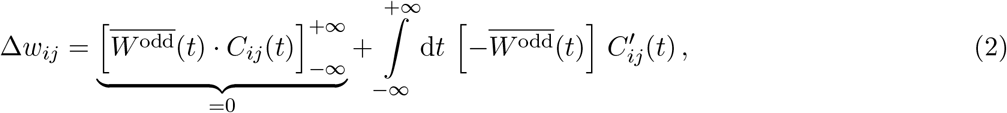

with the primitive 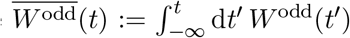 and the derivative 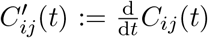. Because 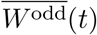 can be assumed to have finite support (note that 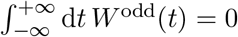), the first term in Equation 2 vanishes. Also the learning window has finite support, and therefore we can restrict the integral in the second term in Equation 2 to a finite region of width *K* around zero:

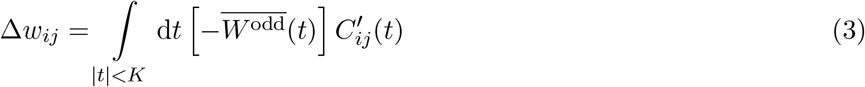

where *K* describes the width of the learning window *W* (gray region in Fig. 2E). The integral in Equation 3 can be interpreted as the cross-correlation’s slope around zero, weighted by the symmetric function 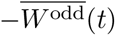 interestingly, features of *C*_*ij*_ for |*t*| ≫ *K*, for example whether side lobes of the correlation function are decreasing or not, are irrelevant.

As a generic example of sequence learning, let us consider the activities of two cells *i* and *j* that encode two behavioral events, for example the traversal of two place fields of two hippocampal place cells. In general, the cells’ responses to these events are called “firing fields”. We model these firing fields as two Gaussian functions *G*_0,*σ*_ and 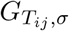 that have the same width *σ* but different mean values 0 and *T*_*ij*_ (we note that *T*_*ij*_ and *σ* are measured in units of time, i.e., seconds; Fig. 2A, dashed curves). In this case of identical Gaussian shapes of the two firing fields, the cross-correlation *C*_*ij*_(*t*) is also a Gaussian function, denoted by 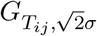, but with mean *T*_*ij*_ and width 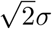 (dashed curve in Fig. 2C). The value *σ* = 0.3 s, which we use in the example of Fig. 2, matches experimental findings on place cells (8, 31).

It is widely assumed that phase precession facilitates temporal-order learning (10, 13, 32), but it has never been quantitatively shown. To test this hypothesis and to calculate how much phase precession contributes to temporal-order learning, we consider Gaussian firing fields that exhibit oscillatory modulations with theta frequency *ω* (Fig. 2A, solid curves). The time-dependent firing rate of cell *i* is described by 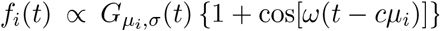, i.e., a Gaussian that is multiplied by a sinusoidal oscillation; see also Equation 11 in *Materials and Methods*. Phase precession occurs with respect to the population theta, which oscillates at a frequency of (1 − *c*)*ω* that is slightly smaller than *ω*, with a “compression factor” *c* that is usually small: 0 ≤ *c* ≪1 (31, 32). This compression factor *c* describes the average advance of the firing phase — from theta cycle to theta cycle — in units of the fraction of a theta cycle; *c* thus determines the slope *ωc* of phase precession (Fig. 2B). A typical value is *c* ≈ *π/*(4*σω*), which accounts for “slope-size matching” of phase precession (31); that is, *c* is inversely proportional to the field size *L* := 4*σ* of the firing field, and the total range of phase precession within the firing field is constant and equals *π* ≡ 180^*?*^. If there are multiple theta oscillation cycles within a firing field (*ωσ* ≫ 1), which is typical for place cells, the cross-correlation *C*_*ij*_(*t*) is a theta modulated Gaussian (solid curve in Fig. 2C; see also Equation 15 in *Materials and Methods*).

The generic shape of the cross-correlation *C*_*ij*_ in Fig. 2C allows for an advanced interpretation of Equation 3, which critically depends on the width *K* of the learning window *W*. We distinguish here two limiting cases: narrow learning windows (*K* ≪1*/ω* ≪*σ*), i.e., the with *K* of the learning window is much smaller than a theta cycle and the width of a firing field, and wide learning windows (*K* ≫ *σ*), i.e., the width *K* of the learning window exceeds the width of a firing field. Let us first consider narrow learning windows. Only later in this manuscript, we will turn to the case of wide learning windows.

### Dependence of temporal-order learning on the overlap of firing fields for narrow learning windows (*K* ≪1*/ω* ≪*σ*)

We first show formally that sequence learning with narrow learning windows requires that the two firing fields do overlap, i.e., their separation *T*_*ij*_ should be less than or at least similar to the width *σ* of the firing fields. In Equation 3, which was derived for odd learning windows, the weight change Δ*w*_*ij*_ is determined by 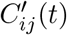 around *t* = 0 in a region of width *K*. For narrow learning windows (*K* ≪1*/ω*), this region is small compared to a theta oscillation cycle and much smaller than the width *σ* of a firing field. Because the envelope of the cross- correlation *C*_*ij*_(*t*) is a Gaussian with mean *T*_*ij*_ and width 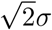, the slope 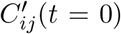 scales with the Gaussian factor 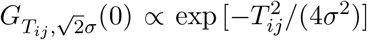. The weight change Δ*w*_*ij*_ therefore strongly depends on the separation *T*_*ij*_ of the firing fields. When the two firing fields do not overlap (*T*_*ij*_ ≫ *σ*), the factor exp 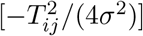 quickly tends to zero, and sequence learning is not possible. On the other hand, when the two firing fields do have considerable overlap (*T*_*ij* ≲_*σ*) we have exp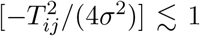 In this case, sequence learning may be feasible with narrow learning windows. In this section, we will proceed the mathematical analysis for overlapping fields, which allows us to assume exp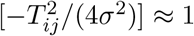.

For overlapping firing fields (*T*_*ij*_ ≲*σ*), let us now consider the fine structure of the cross-correlation *C*_*ij*_(*t*) for |*t*| < *K*, as illustrated in Fig. 2D. Importantly, phase precession causes the first positive peak (i.e., for *t >* 0) of *C*_*ij*_ to occur at time *c T*_*ij*_ with *c* ≪1 (31, 32); phase precession also increases the slope 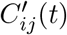 around *t* = 0, which could be beneficial for temporal-order learning according to Equation 3. To quantify this effect, we calculated the cross-correlation’s slope at *t* = 0 (see also Equation 18 in *Materials and Methods*):

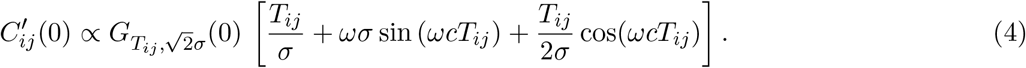

How does 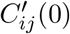 depend on the temporal separation *T*_*ij*_ of the firing fields? If the two fields overlap entirely (*T*_*ij*_ = 0) the sequence has no defined temporal order, and thus 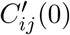 is zero. For at least partly overlapping firing fields (*T*_*ij*_ ≲*σ*) and typical phase precession where *c* = *π/*(4*ωσ*) ≪1, we will show in the next paragraph (and explain in *Materials and Methods* in the text below Equation 18) that the second addend in Equation 4 dominates the other two. In this case, 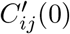 is much higher as compared to the cross-correlation slope in the absence of phase precession (*c* = 0), leading to a clearly larger synaptic weight change for phase precession. The maximum of 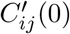 is mainly determined by this second addend (multiplied by 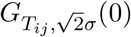) and it can be shown (see *Materials and Methods*) that this maximum is located near 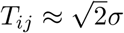.

The increase of 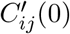 induced by phase precession can be exploited by learning windows *W* that are narrower than a theta cycle (e.g. gray regions in Fig. 2C,D,E). To quantify this effect, let us consider a simple but generic shape of a learning window, for example, the odd STDP window *W* (*t*) = *µ* sign(*t*) exp(−|*t*|*/τ*) with time constant *τ* and learning rate *µ* > 0 (Fig. 2E); this STDP window is narrow for *τ* ≪1*/ω*. Equations 3 and 4 then lead to (see *Materials and Methods*, Equation 19) the average weight change

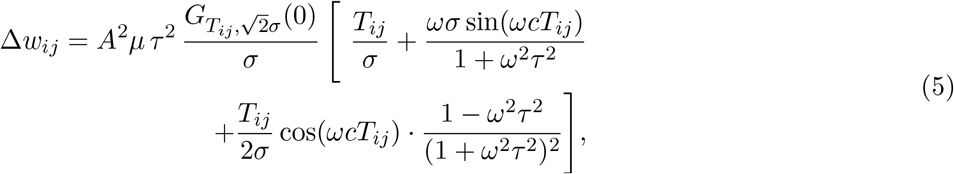

where *A* depicts the number of spikes per field traversal. Note that, according to Equation 3, the weight change Δ*w*_*ij*_ in Equation 5 can be interpreted as a time-averaged version of 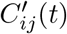 near *t* = 0 from Equation 4. Thus, Equations 4 and 5 have a similar structure, but Equation 5 includes multiple incidences of the term *ω*^2^*τ* ^2^ that account for this averaging. This term is small for narrow learning windows (*τ* ≪1*/ω*) and can thus be neglected (*ω*^2^*τ* ^2^ ≪1) in this limiting case; however, for typical biological values of *τ* ≥ 10 ms and *ω* = 2*π ·* 10 Hz, the peculiar structure of the *ω*^2^*τ* ^2^-containing factor in the third addend in the square brackets is the reason why this addend can be neglected compared to the first one; as a result, the cases of “phase locking” (*c* = 0) and “no theta” (only the first addend remains) are basically indistinguishable. Moreover, for narrow odd learning windows, Δ*w*_*ij*_ in Equation 5 inherits a number of properties from 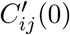 in Equation 4: the second addend still remains the dominant one for *T*_*ij*_ ≲ *σ*; inherited are also the absence of a weight change for fully overlapping fields (Δ*w*_*ij*_ = 0 for *T*_*ij*_ = 0), the maximum weight change for 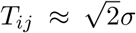, and Δ*w*_*ij*_ → 0 for *T*_*ij*_ → ∞ (Fig. 3A). Furthermore, the prefactor *A*^2^*µτ* ^2^ in Equation 5 suggests that the average weight change increases with increasing width *τ* of the learning window, but we emphasize that this increase is restricted to *τ* ≪1*/ω* (as we assumed for the derivation), which prohibits a generalization of the quadratic scaling to large *τ*; the exact dependence on *τ* will be explained later.

**Figure 3:**
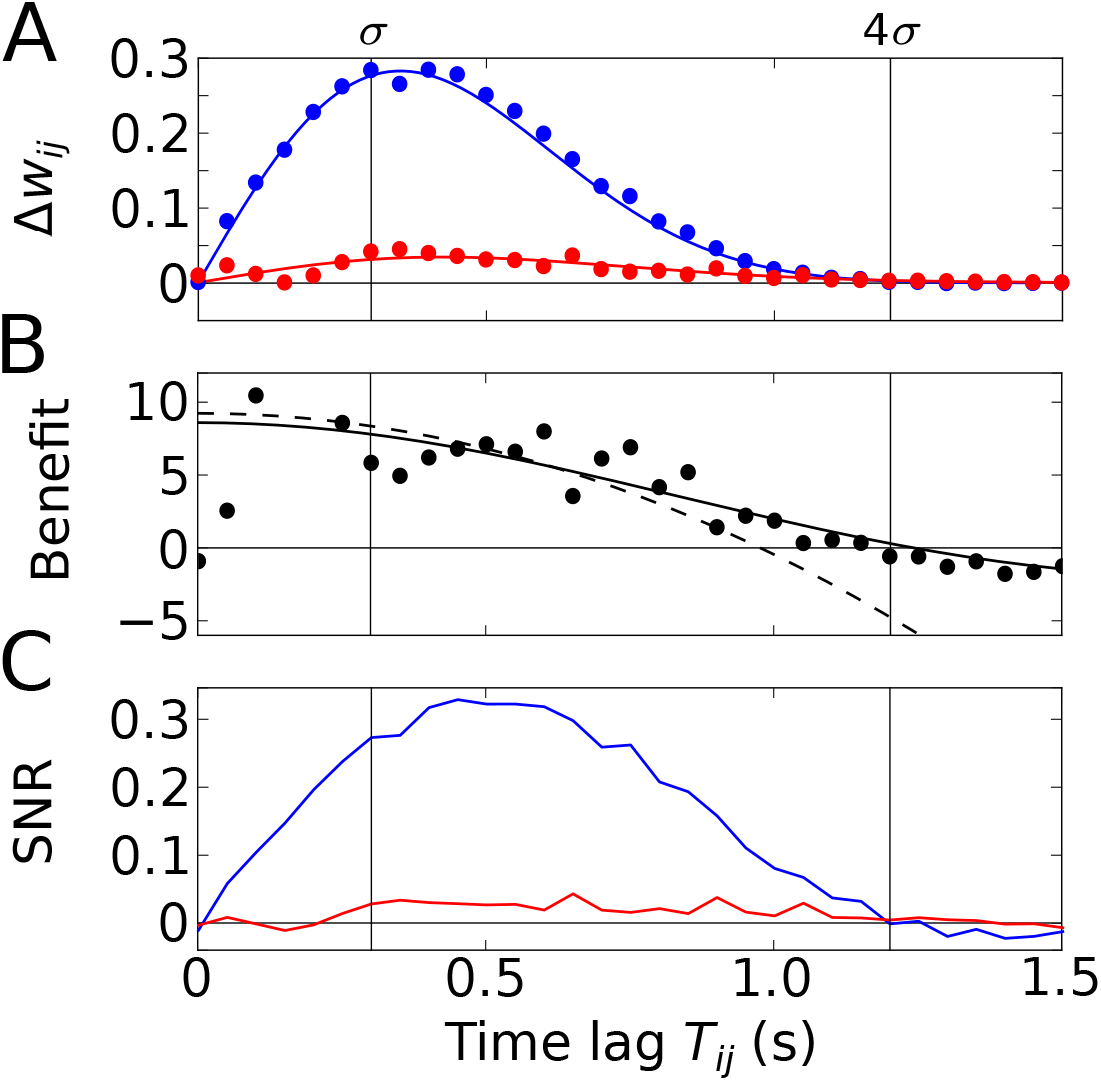
Temporal-order learning for narrow learning windows. (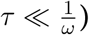. **(A)** The average synaptic weight change Δ*w*_*ij*_ depends on the temporal separation *T*_*ij*_ between the firing fields. Phase precession (blue) yields higher weight changes than phase locking (red). Simulation results (circles, averaged across 10^4^ repetitions) and analytical results (lines, Equation 5) match well. The vertical lines mark time lags of *σ* and 4*σ*, respectively, where 4*σ* approximates the total field width. **(B)** The benefit *B* of phase precession is determined by the ratio of the average weight changes of two scenarios from (A). The solid and dashed lines depict the analytical expression for the benefit (Equation 20) and its approximation for small *T*_*ij*_ (Equation 7), respectively. **(C)** Signal-to-noise ratio (SNR) of the weight change as a function of the firing-field separation *T*_*ij*_. The SNR is defined as the mean weight change divided by the standard deviation across trials in the simulation. Colors as in (A). Parameters for all plots: *ω* = 2 *π ·* 10 Hz, *σ* = 0.3 s, *τ* = 10 ms, *µ* = 1, *c* = 0.042, *A* = 10.

To quantify how much better a sequence can be learned with phase precession as compared to phase locking, we use the ratio of the weight change Δ*w*_*ij*_ with phase precession (*c >* 0) and the weight change Δ*w*_*ij*_(*c* = 0) without phase precession (Fig. 3A), and define the benefit *B* of phase precession as

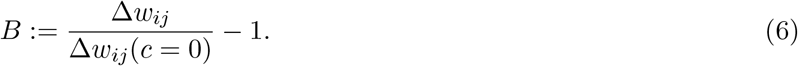

By inserting Equation 5 in Equation 6, we can explicitly calculate the benefit *B* of phase precession (see Equation 20 in *Materials and Methods* and solid line in Fig. 3B). For *T*_*ij*_ ≲ *σ* and *ω*^4^*τ* ^4^ ≪1 (see *Materials and Methods*) the benefit *B* is well approximated by a Taylor expansion up to third order in *T*_*ij*_ (dashed line in Fig. 3B),

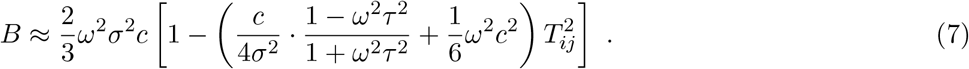

The maximum of *B* as a function of *T*_*ij*_ is obtained for *T*_*ij*_ = 0 (fully overlapping fields), but the average weight change Δ*w*_*ij*_ is zero at this point. We note, however, that *B* decays slowly with increasing *T*_*ij*_, so *B* (*T*_*ij*_ = 0) can be used to approximate the benefit for small field separations *T*_*ij*_ (i.e. largely overlapping fields). For narrow (*ωτ* ≪1) odd STDP windows and slope-size matching (*ωσc* = *π/*4), we find the maximum *B*_max_ ≈ *ωσ/*2, which has an interesting interpretation: If we relate *σ* to the field size *L* of a Gaussian firing field through *L* = 4*σ* and if we relate the frequency *ω* to the period *T*_*θ*_ of a theta oscillation cycle through *T*_*θ*_ = 2*π/ω*, we obtain *B*_max_ ≈ 0.82 *L/T*_*θ*_, i.e., the maximum benefit of phase precession is about the number of theta oscillation cycles in a firing field. The example in Figure 3B (with firing fields in Fig. 2A) has the maximum benefit *B*_max_ ≈ 10 and the benefit remains in this range for partly overlapping firing fields (0 < *T*_*ij*_ ≲ *σ*). We thus conclude that phase precession can boost temporal-order learning by about an order of magnitude for typical cases in which learning windows are narrower than a theta oscillation cycle and overlapping firing fields are an order of magnitude wider than a theta oscillation cycle.

So far, we have considered “average” weight changes that resulted from neural activity that was described by a deterministic firing rate. However, neural activity often shows large variability, i.e., different traversals of the same firing field typically lead to very different spike trains. To account for such variability, we have simulated neural activity as inhomogeneous Poisson processes (see *Materials and Methods* for details). As a result, the change of the weight of a synapse, which depends on the correlation between spikes of the presynaptic and the postsynaptic cells, is a stochastic variable. It is important to consider the variability of the weight change (“noise”) in order to assess the significance of the average weight change. For this reason, we utilize the signal-to-noise ratio (SNR), i.e., the mean weight change divided by its standard deviation (see *Materials and Methods* for details). To do so, we perform stochastic simulations of spiking neurons and calculate the average weight change and its variability across trials. This is done for phase-precessing as well as phase-locked activity. To connect this approach to our previous results, we confirm that the average weight changes estimated from many fields traversals matches well the analytical predictions (Fig. 3A and B, see *Materials and Methods* for details).

The SNR shown in Figure 3C summarizes how reliable is the learning signal in a single traversal of the two firing fields — for the assumed odd learning window. The SNR further depends on *T*_*ij*_ and follows a similar shape as the weight changes in Figure 3A. For phase precession, there is a maximum SNR that is slightly shifted to larger *T*_*ij*_; for phase locking, SNR is always much lower. For the synapse connecting two cells with firing fields as in Figure 2A where *T*_*ij*_ = *σ*, we find a SNR of 0.27, which is insufficient for a reliable representation of a sequence.

To allow reliable temporal-order learning, one possible solution is to increase number of spikes per field traversal *A* (SNR ∝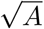, as shown in the Supplemental Information). Another possibility is to increase the number of synapses. In *Materials and Methods* we show that SNR ∝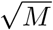 where *M* is the number of identical and uncorrelated synapses. Therefore, to achieve SNR≳ 1 for *A* = 10, one needs *M ≳*14 synapses.

In summary, for narrow, odd learning windows (*τ* ≪1*/ω* ≪*σ*), temporal-order learning could benefit tremendously from phase precession as long as firing fields have some overlap. Average weight changes and the SNR are highest, however, for clearly distinct but still overlapping firing fields. It should be noted that any even component of the learning window would increase the noise and thus further decrease the SNR.

### Dependence of temporal-order learning on the width of the learning window for overlapping firing fields

To investigate how temporal-order learning for an odd learning window depends on its width, we vary the parameter *τ* and quantify the average synaptic weight change Δ*w*_*ij*_ and the SNR both analytically and numerically. We first study overlapping firing fields (Fig. 4) and later consider non-overlapping firing fields (Fig. 5).

**Figure 4:**
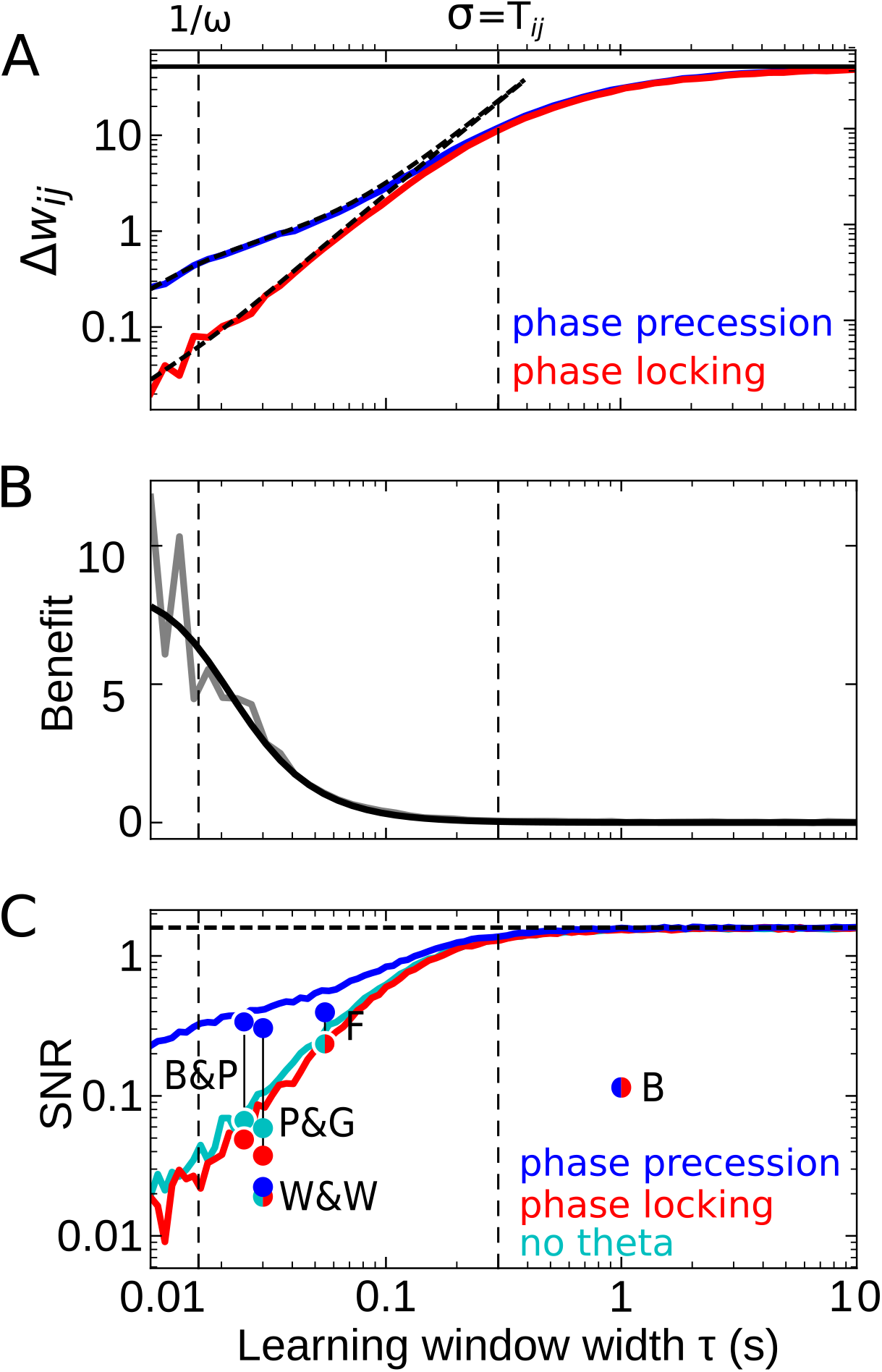
Effect of the learning-window width on temporal-order learning for overlapping fields (here: *T*_*ij*_ = *σ*). **(A)** Average weight change Δ*w*_*ij*_ as a function of width *τ* (for the asymmetric window *W* in Equation 14) for phase precession and phase locking (colored curves). The solid black line depicts the theoretical maximum for large *τ* (Δ*w*_*ij*_ *≈* 52, Equation 8). The dashed curves show the analytical small-tau approximations (Equation 5). The vertical dashed lines mark 1*/ω ≈* 0.016 s and the value of *σ* = *T*_*ij*_ = 0.3 *s*, respectively. **(B)** The benefit *B* of phase precession is largest for narrow learning windows, and it approaches 0 for wide windows. Simulations (gray line) and analytical result (black line, small-tau approximation from Equation 20) match well. **(C)** The signal-to-noise ratio (SNR; phase precession: blue, phase locking: red, no theta: cyan) takes into account that only the asymmetric part of the learning window is helpful for temporal- order learning. For large *τ*, all three coding scenarios induce the same SNR. The horizontal dashed black line depicts the analytical limit of the SNR for large *τ* and overlapping firing fields (SNR *≈* 1.6, Equation 17 of the Supplemental Information). Dots represent the SNR for experimentally observed learning windows. The learning windows were taken from Bi and Poo (“B&P”, ref. 15: their Fig. 1), Froemke et al. (“F”, ref. 16: their Fig. 1D bottom), Wittenberg and Wang (“W&W”, ref. 17: their Fig. 3E), Pfister and Gerstner (“P&G”, ref. 34: their Table 4, ‘All to All’, ‘minimal model’), and Bittner et al. (“B”, ref. 35: their Fig. 3D). For “B&P”, “F”, and “B”, the position of the dots on the horizontal axis was estimated as the average time constants for positive and negative lobes of the learning windows. Wittenberg and Wang modelled their learning rule by a difference of Gaussians — we approximated the corresponding time constant as 30 ms. For the triplet rule by Pfister and Gerstner, we used the average of three time constants: the two pairwise-interaction time constants (as in Bi and Poo) and the triplet-potentiation time constant. Parameters for all plots: *T*_*ij*_ = 0.3 s, *ω* = 2 *π ·* 10 Hz, *σ* = 0.3 s, *c* = 0.042, *A* = 10, *µ* = 1. Colored/gray curves and dots are obtained from stochastic simulations; see *Materials and Methods* for details.

**Figure 5:**
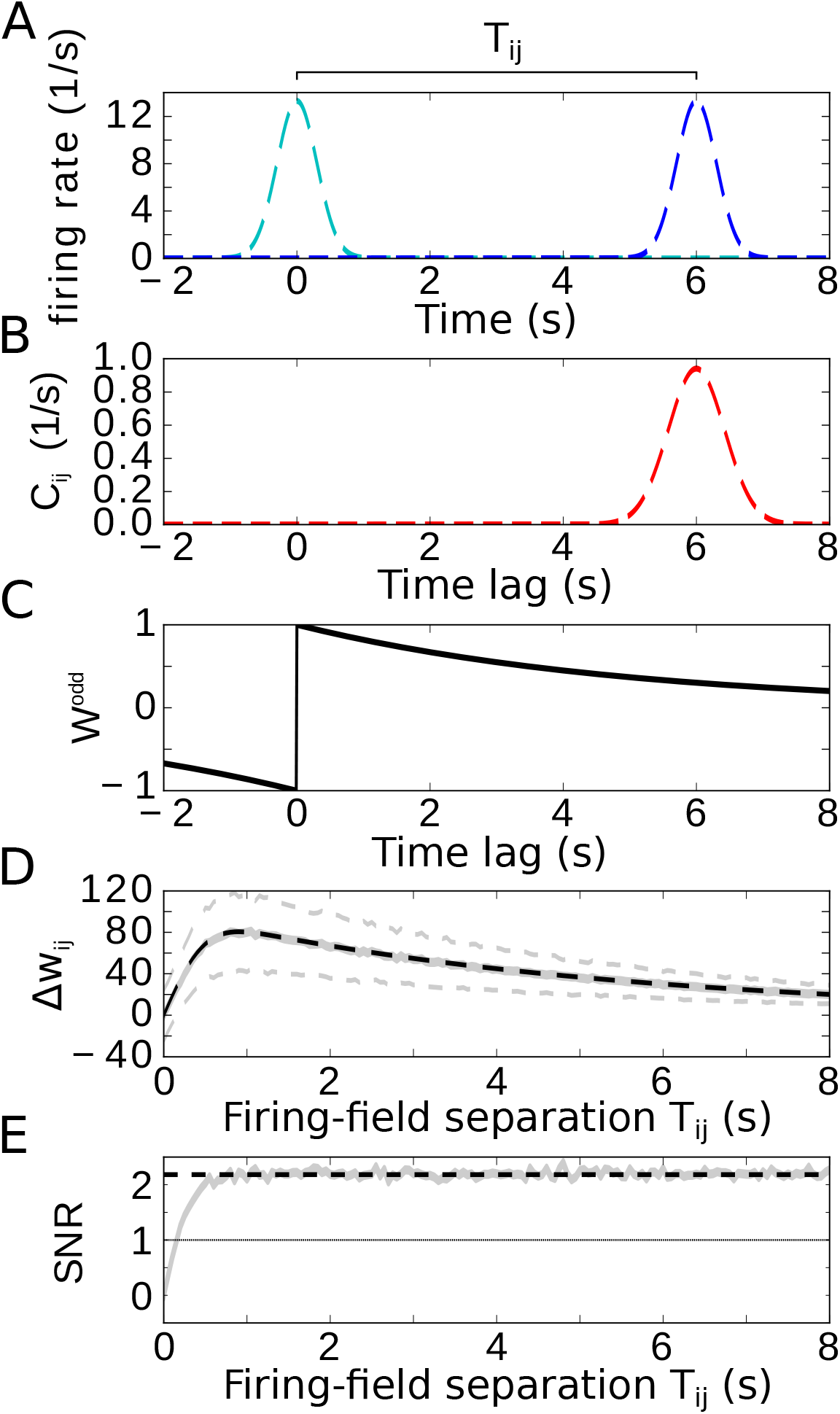
Temporal-order learning for non-overlapping firing fields using wide, asymmetric learning windows. **(A)** Firing rates of two example cells with non-overlapping firing fields. **(B)** Cross-correlation *C*_*ij*_ of the two cells from (A). **(C)** Asymmetric learning window with large width (*τ* = 5s). **(D)** Resulting weight change Δ*w*_*ij*_ for wide learning window and non-overlapping firing fields. The solid grey line depicts the average weight change. The dashed grey lines represent ±1 standard deviation across 1000 repetitions of stochastic spiking simulations. The analytical curve (dashed black line, Equation 9) matches the simulation results. **(E)** SNR of the weight change. Results of the stochastic simulations are shown by the grey curve. The SNR saturates for larger *T*_*ij*_, which fits the analytical expectation (dashed black line, Equation 10). Parameters, unless varied in a plot: *T*_*ij*_ = 6s, *σ* = 0.3 s, *τ* = 5 s, *µ* = 1, *A* = 10.

For partly overlapping firing fields (e.g., *T*_*ij*_ = *σ*), we find numerically that the average synaptic weight change Δ*w*_*ij*_ (the “learning signal”) increases monotonically for increasing *τ* and saturates (colored curves in Fig. 4A). This is because for increasing *τ* the overlap between the learning window and the cross-correlation function grows, and this overlap begins to saturate as soon as the learning window is wider than *T*_*ij*_, i.e., the value at which the cross-correlation assumes its maximum (cmp. dashed curve in Fig. 2C). To analytically calculate the saturation value of Δ*w*_*ij*_ for large learning-window widths (*τ* ≫ *σ*), we can approximate the learning window as a step function (see *Materials and Methods* for details) and find the maximum

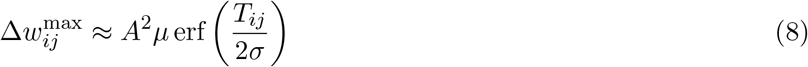

that provides an upper bound to the weight change for overlapping firing fields (solid line in Fig. 4A). For *τ* ≲ 1*/ω* (and actually well beyond this region), the analytical small-tau approximation of Δ*w*_*ij*_ (Equation 5, dashed curves in Fig. 4A) matches the numerical results well.

The results in Figure 4A confirm that Δ*w*_*ij*_ is increased by phase precession for narrow learning windows but is independent of phase precession for *τ* ≫ 1*/ω*. Thus, the benefit *B* becomes small for large *τ* (Fig. 4B) because, for large enough *τ*, the theta oscillation completes multiple cycles within the width of the learning window. To better understand this behavior, let us return to Equation 1: if the product of a *wide* learning window and the cross-correlation *C*_*ij*_ is integrated to obtain the weight change, the oscillatory modulation of the cross-correlation (e.g., as in Fig. 2C) become irrelevant; similarly, according to Equation 3, the particular value of the derivative 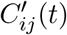 near *t* = 0 can be neglected. Consequently, for *τ* ≫ 1*/ω* phase precession and phase locking as well as the scenario of firing fields that are not theta modulated yield the same weight change (Fig. 4A), and the benefit approaches 0 (Fig. 4B). Wide learning windows thus ignore the temporal (theta) fine-structure of the cross-correlation.

How noisy is this learning signal Δ*w*_*ij*_ across trials? Figure 4C shows that for odd learning windows the SNR increases with increasing *τ* and, for 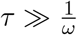, approaches a constant value. This constant value is the same for phase precession, phase locking, or no theta oscillations at all. Taken together, for large enough *τ*, the advantage of phase precession vanishes. For small enough *τ*, phase precession increases the SNR, which confirms and generalizes the results in Fig. 3C. Remarkably, the SNR for “phase locking” is lower than the one for “no theta”, which means that theta oscillations without phase precession degrade temporal-order learning, even though theta oscillations as such were emphasized to improve the modification of synaptic strength in many other cases (e.g. refs. 9, 33).

Figure 4C predicts that a large *τ* yields the biggest SNR, and thus wide learning windows are the best choice for temporal-order learning; however, we note that this conclusion is restricted to odd (i.e., asymmetric) learning windows. An additional even (i.e., symmetric) component of a learning window would increase the noise without affecting the signal, and thus would decrease the SNR (dots in Fig. 4C). It is remarkable that the only experimentally observed instance of a wide window (with *τ* ≈ 1s in ref. 35) has a strong symmetric component, which leads to a low SNR (dot marked “B” in Fig. 4C).

Taken together, we predict that temporal-order learning would strongly benefit from wide, asymmetric windows. However, to date, all experimentally observed (predominantly) asymmetric windows are narrow (e.g. refs. 15–17; see refs. 15, 18 for reviews).

### Temporal-order learning for wide learning windows (*K* ≫ *σ*)

We finally restrict our analysis to wide learning windows, which allows us then to also consider non-overlapping firing fields (Figure 5A,we again use two Gaussians with widths *σ* and separation *T*_*ij*_). To allow for temporal-order learning in this case, the spikes of two non-overlapping fields can only be “paired” by a wide enough learning window. As already indicated in Fig. 4, phase precession does not affect the weight change for such wide learning windows where the width *τ* of the learning window obeys *τ* ≫ 1*/ω* (note that we always assumed many theta oscillation cycles within a firing field, i.e., 1*/ω* ≫ *σ*). Furthermore, Fig. 4 indicated that only the asymmetric part of the learning window contributes to temporal-order learning. For the analysis of temporal-order learning with non-overlapping firing fields and wide learning windows, we thus ignore any theta modulation and phase precession and evaluate, again, only the odd STDP window *W* (*t*) = *µ* sign(*t*) exp(−|*t*|*/τ*). In this case, the weight change (Equation 1) is still determined by the cross-correlation function and the learning window (examples in Fig. 5B, C). The resulting weight change Δ*w*_*ij*_ as a function of the temporal separation *T*_*ij*_ of firing fields is shown in Fig. 5D: with increasing *T*_*ij*_, the weight Δ*w*_*ij*_ quickly increases, reaches a maximum, and slowly decreases. The initial increase is due to the increasing overlap of the Gaussian bump in *C*_*ij*_ with the positive lobe of the learning window. The decrease, on the other hand, is dictated by the time course of the learning window. For *τ* ≫ *σ*, these two effects can be approximated by

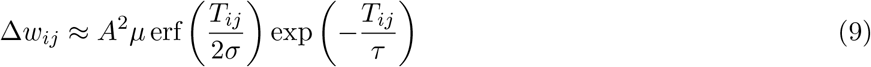

in which the error function describes the overlap of the cross-correlation with the learning window and the exponential term describes the decay of the learning window (dashed black curve in Fig. 5D, see also Equation 25 in *Materials and Methods* for details).

How does the SNR of the weight change depend on the separation *T*_*ij*_ of firing fields? For *T*_*ij*_ = 0, the signal is zero and thus also the SNR. As *T*_*ij*_ increases, both signal and noise increase, but quickly settle on a constant ratio. The value of the SNR height of this plateau can be approximated by

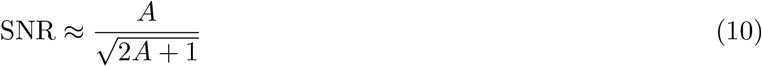

(dashed line in Fig. 5E), where *A* is the number of spikes within a firing field (Equation 11). For *A* = 10, we find SNR ≈ 2.2, allowing for temporal-order learning with a single synapse. We note that this conclusion is limited to asymmetric STDP windows. A symmetric component (like in ref. 35) decreases the SNR and makes temporal-order learning less efficient.

Taken together, temporal-order learning can be performed with wide STDP windows, and phase precession does not provide any benefit; but temporal-order learning requires a purely asymmetric plasticity window. For non-overlapping firing fields, wide learning windows are essential to bridge a temporal gap between the fields.

## Discussion

In this report, we show that phase precession facilitates the learning of the temporal order of behavioral sequences for asymmetric learning windows that are shorter than a theta cycle. To quantify this improvement, we use additive, pairwise STDP and calculate the expected weight change for synapses between two activated cells in a sequence. We confirm the long-held hypothesis (13) that phase precession bridges the vastly different time scales of the slow sequence of behavioral events and the fast STDP rule. Synaptic weight changes can be an order of magnitude higher when phase precession organizes the spiking of multiple cells at the theta time scale as compared to phase-locking cells.

### Other mechanisms and models for sequence learning

As an alternative mechanism to bridge the time scales of behavioral events and the induction of synaptic plasticity, Drew and Abbott (4) suggested STDP and persistent activity of neurons that code for such events. The authors assume regularly firing neurons that slowly decrease their firing rate after the event and show that this leads to a temporal compression of the sequence of behavioral events. For stochastically firing neurons, this approach is similar to ours with two overlapping, unmodulated Gaussian firing fields. In this case, sequence learning is possible, but the efficiency can be improved considerably by phase precession.

Sato and Yamaguchi (20) as well as (22) investigated the memory storage of behavioral sequences using phase precession and STDP in a network model. In computer simulations they find that phase precession facilitates sequence learning, which is in line with our results. In contrast to these approaches, our study focuses on a minimal network (two cells), but this simplification allows us to (i) consider a biologically plausible implementation of STDP, firing fields, and phase precession and (ii) derive analytical results. These mathematical results predict parameter dependencies, which is difficult to achieve with only computer simulations.

Related to our work is also the approach by Masquelier and colleagues (36) who showed that pattern detection can be performed by single neurons using STDP and phase coding, yet they did not include phase precession. They consider patterns in the input whereas, in our framework, it might be argued that patterns between input and output are detected instead.

### Noisy activity of neurons and prediction of the minimum number of synapses for temporal-order learning

To account for stochastic spiking, we use Poisson neurons. We find that a single synapse is not sufficient to reliably encode a minimal two-neuron sequence in a single trial because the fluctuations of the weight change are too large. Fortunately, the SNR scales with 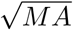, i.e., the square root of the number *M* of identical, but independent synapses and the number *A* of spikes per field traversal of the neurons. For generic hippocampal place fields and typical STDP, we predict that about 14 synapses are sufficient to reliably encode temporal order in a single traversal. Interestingly, peak firing rates of place fields are remarkably high (up to 50 spikes/s; e.g. 8, 37). Taken together, in hippocampal networks, reliable encoding of the temporal order of a sequence is possible with a low number of synapses, which matches simulation results on memory replay (24).

### Width, shape, and symmetry of the STDP window are critical for temporal-order learning

Various widths have been observed for STDP learning windows (15, 18). We show that for all experimentally found STDP time constants phase precession can improve temporal-order learning. However, for learning windows much wider than a theta oscillation cycle, the benefit of phase precession for temporal-order learning is small. Wide learning windows, where the width can be even on a behavioral time scale of ≈ 1s (35) or larger, could, on the other hand, enable the association of non-overlapping firing fields. Alternatively, non-overlapping firing fields might also be associated by narrow learning windows if additional cells (with firing fields that fill the temporal gap) help to bridge a large temporal difference, much like “time cells” in the hippocampal formation (reviewed in 38).

STDP windows typically have symmetric and asymmetric components (18). We find that only the asymmetric component supports the learning of temporal order. In contrast, the symmetric component strengthens both forward and backward synapses by the same amount and thus contributes to the association of behavioral events independent of their temporal order. For example, the learning window reported by Bittner et al. (35) shows only a mild asymmetry and is thus unfavorable to store the temporal order of behavioral events. Only long, predominantly asymmetric STDP windows would allow for effective temporal-order learning (cf. Fig. 4).

Generally, the shape of STDP windows is subject to neuromodulation; for example, cholinergic and adrenergic modulation can alter its polarity and symmetry (39). Also dopamine can change the symmetry of the learning window (40). Therefore, sequence learning could be modulated by the behavioral state (attention, reward, etc.) of the animal.

### Key features of phase precession for temporal order-learning: generalization to non-periodic modulation of activity

For STDP windows narrower (≲ 10 ms) than a theta cycle (≳ 100 ms), we argue that the slope of the cross-correlation function at zero offset controls the change of the weight of the synapse connecting two neurons; and we show that phase precession can substantially increase this slope. This result predicts that features of the cross-correlation at temporal offsets that are larger than the width of the learning window are irrelevant for temporal-order learning. It is thus conceivable to boost temporal-order learning even without phase precession, which is weak if theta oscillations are weak, as for example in bats (41) and humans (42). In this case, temporal-order learning may instead benefit from two other phenomena that could create an appropriate shape of the cross-correlation: (1) Spiking of cells is locked to common (aperiodic) fluctuations of excitability. (2) Each cell responds the faster to an increase in its excitability the longer ago its firing field has been entered, which may be mediated by a progressive facilitation mechanism. Together, these phenomena can make the cross-correlation exhibit a steeper slope around zero and could even give rise to a local maximum at a positive offset. This temporal fine structure is superimposed on a slower modulation, which is related to the widths of the firing fields. In summary, a progressively decreasing delay of spiking with respect to non-rhythmic fluctuations in excitation generalizes the notion of phase precession. Interestingly, synaptic short-term facilitation, which could generate the described fine structure of the cross-correlation, has also been proposed as mechanism underlying phase precession (43).

### Model assumptions

In our model we assumed that recurrent synapses (e.g. between neurons representing a sequence) are plastic but weak during encoding, such that they have a negligible influence on the postsynaptic firing rate; and that the feedforward input dominates neuronal activity. These assumptions seem justified as Hasselmo (39) indicated that excitatory feedback connections may be suppressed during encoding to avoid interference from previously stored information (see also 44). Furthermore, neuromodulators facilitate long-term plasticity (reviewed, e.g., by 45), which also supports our assumptions.

The assumption of weak recurrent connections implies that these connections do not affect the dynamics. Consequently (and in contrast to ref. 14), we thus hypothesize that phase precession is not generated by the local, recurrent network (see also, e.g. 46); instead, we assume that phase precession is inherited from upstream feedforward inputs (47, 48) or generated locally by a cellular/synaptic mechanism (49–52). After temporal-order learning was successful, the resulting asymmetric connections could indeed also generate phase precession (as demonstrated by the simulations in ref. 14), and this phase precession could then even be similar to the one that has initially helped to shape synaptic connections. Finally, inherited or local cellularly/synaptically-generated phase precession and locally network-generated phase precession could interact (as reviewed, for example in ref. 53).

We assumed in our model that the widths of the two firing fields that represent two events in a sequence are identical (see, e.g., Fig. 2A). But firing fields may have different widths, and in this case a slope-size matched phase precession would fail to reproduce the timing of spikes required for the learning of the correct temporal order of the two events. For example, the learned temporal order of events (timed according to field entry) would even be reversed if two fields with different sizes are aligned at their ends. How could the correct temporal order nevertheless be learned in our framework? In the hippocampus, theta oscillations are a traveling wave (54, 55) such that there is a positive phase offset of theta oscillations for the wider firing fields in the more ventral parts of the hippocampus. This traveling-wave phenomenon could preserve the temporal order in the phase-precession-induced compressed spike timing, as also pointed out earlier (56, 57).

Our results on learning rules for sequence learning rely on pairwise STDP in which pairs of presynaptic and postsynaptic spikes are considered. Conversely, triplet STDP considers also motifs of three spikes (either 2 presynaptic - 1 postsynaptic or 2 postsynaptic - 1 presynaptic) (34). Triplets STDP models can reproduce a number of experimental findings that pairwise STDP could not, for example the dependence on the repetition frequency of spike pairs (58). To investigate the influence of triplet interactions on sequence learning, we implemented the generic triplet rule by Pfister and Gerstner (34). We used their “minimal” model, which was regarded as the best model in terms of number of free parameters and fitting error; for the parameters they obtained from fitting the triplet STDP model to hippocampal data, we found only mild differences to our results (see, e.g., Fig. 4C). Differences are small because the fitted time constant of the triplet term (40 ms) is smaller than typical inter-spike intervals (≳ 50 ms, minimum in field centers) in our simulations.

### Replay of sequences and storage of multiple and overlapping sequences

A sequence imprinted in recurrent synaptic weights can be replayed during rest or sleep (27, 59–62), which was also observed in network-simulation studies (25, 26, 63). Replay could thus be a possible readout of the temporal-order learning mechanism. However, replay depends on the many parameters of the network, and a thorough investigation of is beyond the scope of this manuscript. Therefore, we focus on synaptic weight changes that represent the formation of sequences in the network, which underlies replay, and we do not simulate replay.

We have considered the minimal example of a sequence of two neurons. Sequences can contain many more neurons, and the question arises how two different sequences can be told apart if they both contain a certain neuron, but proceed in different directions — as they might do for sequences of spatial or non-spatial events (64). In this case, it may be beneficial to not only strengthen synapses that connect direct successors in the sequence but also synapses that connect the second-to-next neuron. In this way, the two crossing sequences could be disambiguated, and the wider context in which an event is embedded becomes associated, which is in line with retrieved-context theories of serial-order memory (65). More generally, it is an interesting question of how many sequences can be stored in a network of a given size. Gillett et al. (26) were able to analytically calculate the storage capacity for the storage of sequences in a Hebbian network.

In conclusion, our model predicts that phase precession enables efficient and robust temporal-order learning. To test this hypothesis, we suggest experiments that modulate the shape of the STDP window or selectively manipulate phase precession and evaluate memory of temporal order.

## Materials and Methods

### Experimental Design: Model description

We model the time-dependent firing rate of a phase precessing cell *i* (two examples in Fig. 2A) as

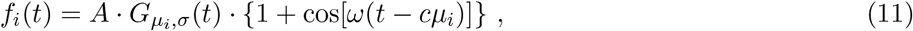

where the scaling factor *A* determines the number of spikes per field traversal and 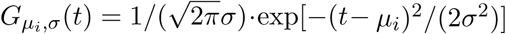 is a Gaussian function that describes a firing field with center at *µ*_*i*_ and width *σ*. The firing field is sinusoidally modulated with theta frequency *ω* (but the sinusoidal modulation is not a critical assumption, see *Discussion*), with typically many oscillation cycles in a firing field (*ωσ* ≫ 1). The compression factor *c* can be used to vary between phase precession (*c >* 0), phase locking (*c* = 0), and phase recession (*c* < 0) because the average population activity of many such cells oscillates at frequency of (1 − *c*)*ω* (31, 33), which provides a reference frame to assign theta phases (Fig. 2A). Usually, |*c*| ≪1 with typical values *c* ≲ 1*/*(*σω*) (31); for a pair of cells with overlapping firing fields (centers separated by *T*_*ij*_ := *µ*_*j*_ − *µ*_*i*_) the phase delay is *ωcT*_*ij*_ (Fig. 2B).

To quantify temporal-order learning, we consider the average weight change Δ*w*_*ij*_ of the synapse from cell *i* to cell *j*, which is (30)

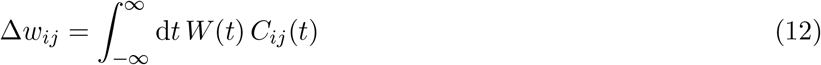

where *C*_*ij*_ (*t*) is the cross-correlation between the firing rates *f*_*i*_ and *f*_*j*_ of cells *i* and *j*, respectively (Fig. 2C,D):

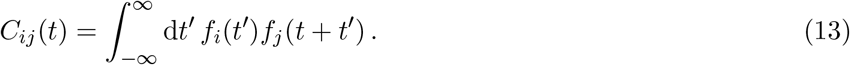

*W* (*t*) denotes the synaptic learning window, for example the asymmetric window

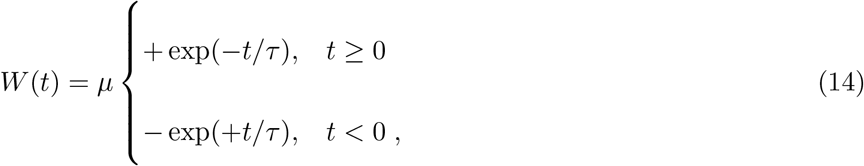

where *τ* is the time constant and *µ >* 0 is the learning rate (Fig. 2E).

For the following calculations, we make two assumptions that are reasonable in the hippocampal formation (8, 15, 31):

i. The theta oscillation has multiple cycles within the Gaussian envelope of the firing field in Equation 11 (1*/ω* ≪*σ*).
ii. The window *W* is short compared to the theta period (*τ* ≪1*/ω*).

### Analytical approximation of the cross-correlation function

To explicitly calculate the cross-correlation *C*_*ij*_(*t*) as defined in Equation 13, we plug in the firing-rate functions (Equation 11) for the two neurons:

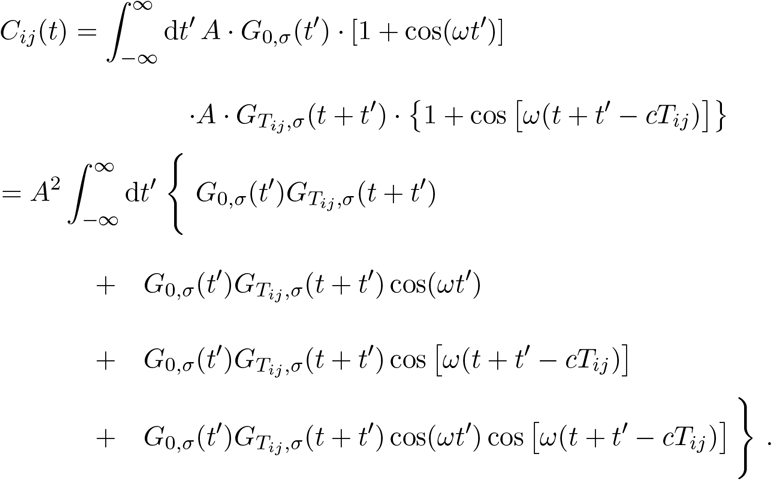

The first term (out of four) describes the cross-correlation of two Gaussians, which results in a Gaussian function centered at *T*_*ij*_ and with width *σ* 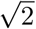. For the second term, we note that the product of two Gaussians yields a function proportional to a Gaussian with width *σ*/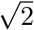, and then use assumption (i). When integrated, the second term’s contribution to *C*_*ij*_(*t*) is negligible because the cosine function oscillates multiple times within the Gaussian bump, i.e., positive and negative contributions to the integral approximately cancel. The same argument applies to the third term. For the fourth term, we use the trigonometric property cos(*α*) *·*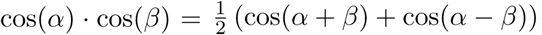. We set *α* = *ωt*′, *β* = *ω*(*t* + *t*′ − *cT*_*ij*_) and find

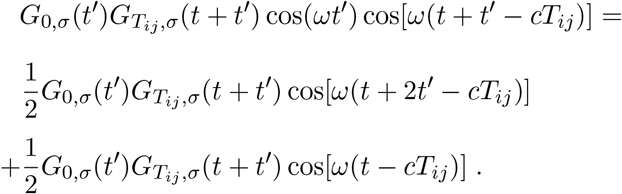

Again, we use assumption (i) and neglect the first addend on the right-hand side. Notably, the cosine function in the second addend is independent of the integration variable *t*. Taken together, we find

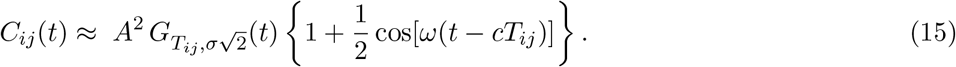

Thus, the cross-correlation can be approximated by a Gaussian function (center at *T*_*ij*_, width *σ* 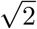) that is theta modulated with an amplitude scaled by the factor 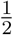.

To further simplify Equation 15, we note that the time constant *τ* of the STDP window is usually small compared to the theta period (assumption (ii), Fig. 2C,D,E). Structures in *C*_*ij*_(*t*) for |*t*| ≫ *τ* thus have a negligible effect on the synaptic weight change. Therefore, we can focus on the cross-correlation for small temporal lags. In this range, we approximate the (slow) Gaussian modulation of *C*_*ij*_(*t*) (Fig. 2C,D, dashed red line) by a linear function, that is,

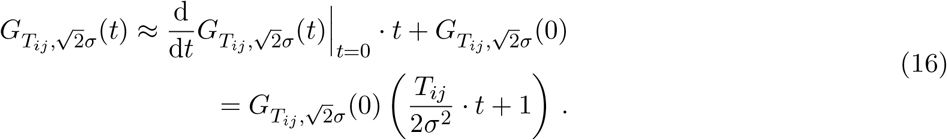

Inserting this result in Equation 15, we approximate the cross-correlation function *C*_*ij*_(*t*) for |*t*| ≲ *τ* as (Fig. 2D, dashed black line)

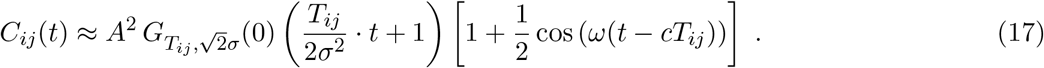

In the *Results* we show that the slope of the cross-correlation function at *t* = 0 is important for temporal-order learning. From Equation 17 we find

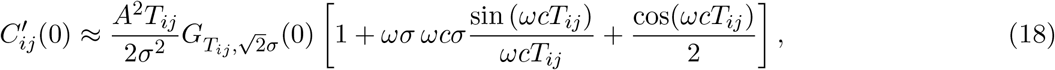

which has three addends within the square brackets. Let us estimate the relative size of the second and third terms with respect to the first one. The third term is at most of the order of 0.5 because | cos(*ωcT*_*ij*_)| ≤ 1. For the second addend, we note that sin(*ωcT*_*ij*_)*/*(*ωcT*_*ij*_) approaches 1 for *T*_*ij*_ *→* 0 and remains in this range for |*ωcT*_*ij*_|≲ *π/*4. This condition is fulfilled for |*T*_*ij*_|≲ *σ* if we assume slope-size matching of phase precession (31), i.e., 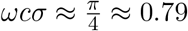. Then, the size of the second addend is dictated by the factor *ωσ*, which is large according to assumption (i). In other words, for typical phase precession and |*T*_*ij*_|≲ *σ*, the second addend is much larger than the other two.

To further understand the structure of 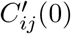 which is also shaped by the prefactors in front of the square brackets, we first note that 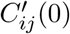 is zero for fully overlapping firing fields (*T*_*ij*_ *→* 0). On the other hand, for very large field separations (*T*_*ij*_ ≫ *σ*), the Gaussian term *G* causes 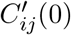 to become zero. The prefactors have a maximum at 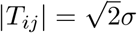. The maximum’s exact location is slightly shifted by the second addend but remains near 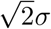. This peak will be important because it is inherited by the average weight change (Equation 3).

### Average weight change

Having approximated the cross-correlation function and its slope at zero (Equations 17,18), we are now ready to calculate the average synaptic weight change (Equation 3) for the assumed STDP window (Equation 14). Standard integration methods yield

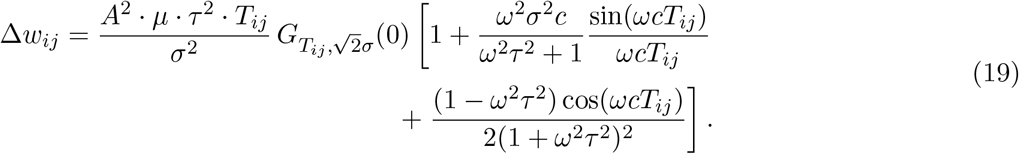

Because Δ*w*_*ij*_ is a temporal average of 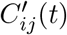 for small *t* (see interpretation of Equation 3), the weight change’s structure resembles the previously discussed structure of 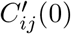. The averaging introduces additional factors proportional to 1 ± *ω*^2^*τ* ^2^, but for *ωτ* ≪1 [assumption (ii)] those have only minor effects on the relative size of the three addends. The second term still dominates. Importantly, Δ*w*_*ij*_ = 0 for *T*_*ij*_ = 0 and the position of the peak at 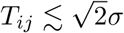 is inherited from 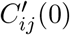 (cf. Fig. 3A).

### The benefit of phase precession

To quantify the benefit *B* of phase precession, we consider the expression Δ*w*_*ij*_*/*Δ*w*_*ij*_(*c* = 0) −1, because Δ*w*_*ij*_ describes the overall weight change (including phase precession), and Δ*w*_*ij*_(*c* = 0) serves as the baseline weight change due to the temporal separation of the firing fields (without phase precession). We subtract 1 to obtain *B* = 0 when the weight changes are the same with and without phase precession. From Equation 19 we find

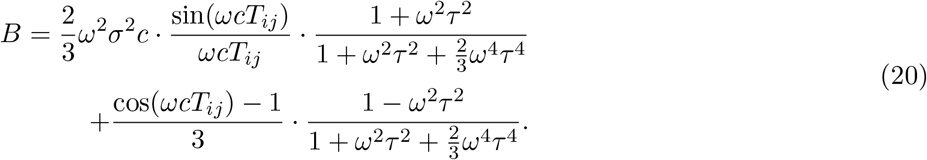

To better understand the structure of *B*, we Taylor-expand it in *T*_*ij*_ up to the third order and assume *ω*^4^*τ* ^4^ ≪1 [cf. assumption (ii)]. The result is

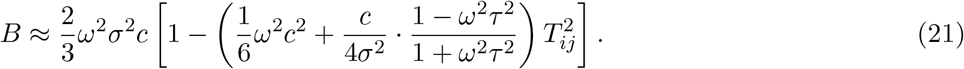

Thus, *B* assumes a maximum for *T*_*ij*_ = 0 and slowly decays for small *T*_*ij*_ (Fig. 3B). Using slope-size matching (*ωσc* = *π/*4), the maximal benefit is

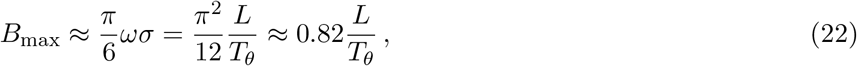

where *L* = 4*σ* depicts the total field size and 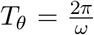 is the period of the theta oscillation. Thus, the number of theta cycles per firing field determines the benefit for small separations of the firing fields.

### Average weight change for wide learning windows

In this paragraph we relax assumption (ii), i.e., we consider wide asymmetric learning windows *W* (Equation 14 with *τ* ≫ *σ*). Furthermore, we neglect any theta-oscillatory modulation of the firing fields in Equation 11 and, thus, *C*_*ij*_ in Equation 15.

First, for non-overlapping fields (*T*_*ij*_ ≫ *σ*), the learning window can be approximated to be constant near the peak of the Gaussian bump of *C*_*ij*_. We can thus rewrite Equation 1 as

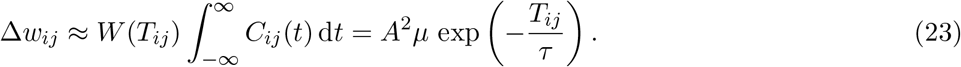

Second, for overlapping fields (0 < *T*_*ij*_ ≲*σ*), the Gaussian bump of *C*_*ij*_ partly lies on the negative lobe of *W*. We can approximate *W* (*t*) = sign(*t*), and the average weight change in Equation 1 then reads

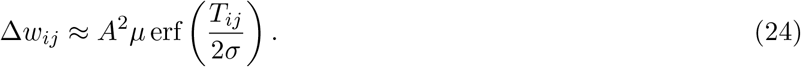

Combining the two limiting cases in Equations 23 and 24 yields

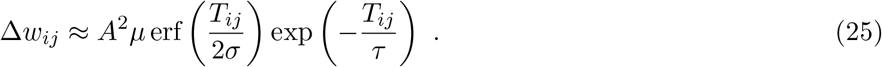

### Signal-to-noise ratio

To correctly encode the temporal order of behavioral events, the average weight change Δ*w*_*ij*_ of a forward synapse needs to be larger than the average weight change Δ*w*_*ji*_ of the corresponding backward synapse. We thus define the signal-to-noise ratio as

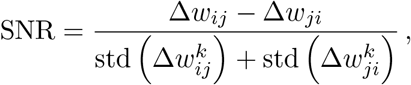

where std() denotes the standard deviation and 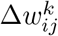,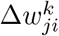 are the weight changes for trial *k* ∈ [1, *N*], the averages across trials being 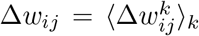 and 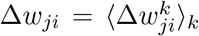. This expression for the SNR “punishes” the non-sequence-specific strengthening of backward synapses. Specifically, SNR = 0 for a symmetric (even) learning window, because the numerator (which represents the “signal”) is zero. On the other hand, a perfectly asymmetric learning window, like the one used throughout this study (Equation 14), yields SNR 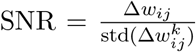, because 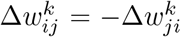. Asymmetric learning windows thus recover the classical definition of the SNR as the ratio between the average weight change and the standard deviation of the weight change.

We note that the generalized definition above can be used to calculate the SNR for arbitrary windows, such as the learning window from Bittner et al. (2017, Figure 4C).

Assuming an asymmetric window and *M* uncorrelated synapses with the same mean and variance of the weight change, we can write the signal-to-noise ratio as

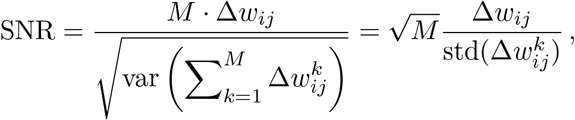

because the variance of the sum can be decomposed into the sum of variances and covariances. All covariances are zero because synapses are uncorrelated. This leaves a sum of *M* variances, which are identical. Therefore, the standard deviation, and consequently also the SNR, scale with 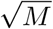.

### Numerical simulations

To numerically simulate the synaptic weight change, spikes were generated by inhomogeneous Poisson processes with rate functions according to Equation 11. For every spike pair, the contribution to the weight change was calculated according to Equation 14. We repeated the simulations for *N* = 10^4^ trials, and the mean weight change as well as the standard deviation across trials and the SNR were estimated.

## Supporting information

Supplemental mathematical derivations

## Acknowledgements

This work was funded by the Deutsche Forschungsgemeinschaft (DFG, German Research Foundation; Grants GRK 1589/2, SPP 1665, SFB 1315 – project-ID 327654276) and the German Federal Ministry for Education and Research (BMBF; Grant 01GQ1705). We thank Ikhwan Khalid, Lukas Kunz, Natalie Schieferstein, Tiziano D’Albis, Paul Pfeiffer, and Adam Wilkins for helpful discussions and feedback on the manuscript. ETR and RK designed and performed the research, wrote and discussed the manuscript. ETR and RK declare no conflict of interest.

