## Supplemental mathematical derivations for "Synaptic learning rules for sequence learning"

### Supplementary Information

#### The signal-to-noise ratio of $\Delta w_{ij}$

A synapse with weight $w_{ij}$ is assumed to connect neuron $i$ to neuron $j$. Here, we aim to derive the signal-to-noise ratio (SNR) of the weight changes $\Delta w_{ij}$, which is defined as (*Materials and Methods*)

$$\mathrm{SNR} = \frac{\langle \Delta w_{ij} \rangle - \langle \Delta w_{ji} \rangle}{\mathrm{std}\,(\Delta w_{ij}) + \mathrm{std}\,(\Delta w_{ji})} \,. \tag{1}$$

where $\langle \Delta w_{ij} \rangle$ is the average signal. The noise is described by the standard deviation of the weight change,

$$\mathrm{std}\,(\Delta w_{ij}) = \sqrt{\mathrm{var}(\Delta w_{ij})} = \sqrt{\langle \Delta w_{ij}^2 \rangle - \langle \Delta w_{ij} \rangle^2} \,.$$

Signal and noise are generated by additive STDP and spiking activity that is modelled by two inhomogeneous Poisson processes with rates $f_i(t)$ and $f_j(t)$ that have finite support. The average weight change is calculated by $\langle \Delta w_{ij} \rangle = \int \mathrm{d}t\, W(t) C_{ij}(t)$ where $W(t)$ is the synaptic learning window and $C_{ij}(t)$ depicts the cross-correlation function $C_{ij}(t) = \int \mathrm{d}t' f_i(t') f_j(t'+t)$. From Kempter et al. (1999), we use

$$\begin{aligned}
\langle \Delta w_{ij}^2 \rangle(t) = -\Delta w_{ij}(t_0)^2 + 2\Delta w_{ij}(t_0)\langle \Delta w_{ij} \rangle(t) + \int_{t_0}^{t} \mathrm{d}t' \int_{t_0}^{t} \mathrm{d}u \\
\Bigg\{ \langle S_i(t') S_i(u) \rangle (w^{\mathrm{in}})^2 + \langle S_j(t') S_j(u) \rangle (w^{\mathrm{out}})^2 + \langle S_i(t') S_j(u) \rangle 2 w^{\mathrm{in}} w^{\mathrm{out}} + \\
2 \int \mathrm{d}s\, W(s) \left( \langle S_i(t') S_i(u+s) S_j(u) \rangle w^{\mathrm{in}} + \langle S_j(t') S_i(u+s) S_j(u) \rangle w^{\mathrm{out}} \right) + \\
\int \mathrm{d}s \int \mathrm{d}v\, W(s) W(v) \langle S_i(t'+s) S_i(u+v) S_j(t') S_j(u) \rangle \Bigg\} \,,
\end{aligned} \tag{2}$$

where $S_i(t) = \sum_n \delta(t - t_i^{(n)})$ and $S_j(t) = \sum_n \delta(t - t_j^{(n)})$ are the presynaptic and postsynaptic spike trains, respectively. To simplify, we set $t_0 = -\infty$ and $\Delta w_{ij}(t_0) = 0$. Furthermore, we are interested in paired STDP and thus set $w^{\mathrm{in}} = w^{\mathrm{out}} = 0$. For $\langle \Delta w_{ij}^2 \rangle = \lim_{t \to \infty} \langle \Delta w_{ij}^2 \rangle(t)$, Equation 2 reduces to

$$\langle \Delta w_{ij}^2 \rangle = \int_{-\infty}^{\infty} \mathrm{d}t' \int_{-\infty}^{\infty} \mathrm{d}u \int \mathrm{d}s \int \mathrm{d}v\, W(s) W(v) \langle S_i(t'+s) S_i(u+v) S_j(t') S_j(u) \rangle . \tag{3}$$

Because both spike trains are drawn from different Poisson processes, $S_i$ and $S_j$ are statistically independent, and therefore we can simplify

$$\langle S_i(t'+s)S_i(u+v)S_j(t')S_j(u)\rangle = \langle S_i(t'+s)S_i(u+v)\rangle\langle S_j(t')S_j(u)\rangle.$$

Moreover, in a spike train the spikes at different times are uncorrelated,

$$\langle S_i(t'+s)S_i(u+v)\rangle = \langle S_i(t'+s)\rangle\langle S_i(u+v)\rangle + \langle S_i(t'+s)\rangle\delta(t'+s-u-v)$$

and

$$\langle S_j(t')S_j(u)\rangle = \langle S_j(t')\rangle\langle S_j(u)\rangle + \langle S_j(t')\rangle\delta(t'-u).$$

As $S_i$ and $S_j$ are realizations of inhomogeneous Poisson processes with rates $f_i(t)$ and $f_j(t)$, respectively, we find

$$\langle S_i(t'+s)S_i(u+v)\rangle = f_i(t'+s)f_i(u+v) + f_i(t'+s)\delta(t'+s-u-v)$$

and

$$\langle S_j(t')S_j(u)\rangle = f_j(t')f_j(u) + f_j(t')\delta(t'-u).$$

We insert these expressions into Equation 3:

$$\langle \Delta w_{ij}^2\rangle = \int_{-\infty}^{\infty} \mathrm{d}t' \int_{-\infty}^{\infty} \mathrm{d}u \int \mathrm{d}s \int \mathrm{d}v\, W(s)W(v)F(t',s,u,v) \tag{4}$$

where

$$\begin{aligned} F(t',s,u,v) = &[f_i(t'+s)f_i(u+v) + f_i(t'+s)\delta(t'+s-u-v)] \\ &\cdot [f_j(t')f_j(u) + f_j(t')\delta(t'-u)]\,. \end{aligned}$$

To explicitly calculate the SNR, we parameterize the firing rates as

$$f_i(t) = \frac{A}{\sqrt{2\pi}\sigma}\exp\left(-\frac{t^2}{2\sigma^2}\right)\cdot[1+\cos(\omega t)] \tag{5}$$

and

$$f_j(t) = \frac{A}{\sqrt{2\pi}\sigma}\exp\left(-\frac{(t-T_{ij})^2}{2\sigma^2}\right)\cdot[1+\cos(\omega(t-cT_{ij}))]\,. \tag{6}$$

See main text for definitions of symbols. Furthermore, we assume $W(s) = W^{\text{odd}}(s) + W^{\text{even}}(s)$ with

$$W^{\text{odd}}(s) = \mu\begin{cases} +\exp(-s/\tau), & s \geq 0 \\ -\exp(+s/\tau), & s < 0\,, \end{cases}$$

(see Equation 14 in *Materials and Methods*) and

$$W^{\text{even}} = \lambda\exp(-|s|/\kappa)\,.$$

In what follows we consider a limiting case of wide learning windows, for which we can explicitly calculate the SNR. The results obtained in this case match well to the numerical simulations for wide learning windows (Figs. 4 and 5 in the main text).

#### Wide learning windows

For wide windows (formally: $\tau \to \infty$, $\kappa \to \infty$), we can approximate $W^{\text{even}} = \lambda$ and $W^{\text{odd}}(t) = \mu \operatorname{sgn}(t)$ and neglect the sinusoidal modulations of $f_i$ and $f_j$ in Equations 5 and 6; phase precession does not affect the SNR in this case.

The following calculations are similar for odd and even windows. We elaborate the calculations in detail for odd windows and use '$\pm$' and '$\mp$' to include the similar calculations for even windows. The top symbol ('+' and '-', respectively) corresponds to odd windows; the bottom symbol corresponds to even windows.

To start, we split the third and fourth integral in Equation 4 into positive and negative time lags $s$ and $v$, respectively:

$$\begin{aligned}\frac{1}{\mu^2}\langle \Delta w_{ij}^2 \rangle(t) &= \int_{-\infty}^{\infty} \mathrm{d}t' \int_{-\infty}^{\infty} \mathrm{d}u \int \mathrm{d}s \int \mathrm{d}v \operatorname{sgn}(s) \operatorname{sgn}(v) F(t', s, u, v) \\ = \int_{-\infty}^{\infty} \mathrm{d}t' \int_{-\infty}^{\infty} \mathrm{d}u \Bigg\{ & \int_{0}^{\infty} \mathrm{d}s \int_{0}^{\infty} \mathrm{d}v F(t', s, u, v) \\ & \mp \int_{0}^{\infty} \mathrm{d}s \int_{-\infty}^{0} \mathrm{d}v F(t', s, u, v) \\ & \mp \int_{-\infty}^{0} \mathrm{d}s \int_{0}^{\infty} \mathrm{d}v F(t', s, u, v) \\ & + \int_{-\infty}^{0} \mathrm{d}s \int_{-\infty}^{0} \mathrm{d}v F(t', s, u, v) \Bigg\}\end{aligned} \tag{7}$$

We rewrite $F$ as

$$\begin{aligned}F(t', s, u, v) = &\overbrace{f_i(t'+s) f_i(u+v) f_j(t') f_j(u)}^{\text{(i)}} + \overbrace{f_i(t'+s) f_i(u+v) f_j(t') \delta(t'-u)}^{\text{(ii)}} + \\ &\underbrace{f_i(t'+s) \delta(t'+s-u-v) f_j(t') f_j(u)}_{\text{(iii)}} + \underbrace{f_i(t'+s) \delta(t'+s-u-v) f_j(t') \delta(t'-u)}_{\text{(iv)}},\end{aligned}$$

which has four addends and occurs in four integrals in Equation 7. Thus, there are 16 terms we need to evaluate. We label these terms (1.i) to (1.iv) for the first integral, (2.i) to (2.iv) for the second integral and so on until (4.iv).

For the term (1.i) we find

$$
\begin{aligned}
&\int_{-\infty}^{\infty}\mathrm{d}t'\int_{-\infty}^{\infty}\mathrm{d}u\int_{0}^{\infty}\mathrm{d}s\int_{0}^{\infty}\mathrm{d}v f_i(t'+s)f_i(u+v)f_j(t')f_j(u)\\
=&\int_{-\infty}^{\infty}\mathrm{d}t' f_j(t')\int_{-\infty}^{\infty}\mathrm{d}u f_j(u)\int_{0}^{\infty}\mathrm{d}s f_i(t'+s)\int_{0}^{\infty}\mathrm{d}v f_i(u+v)\\
=&\frac{A^2}{4}\int_{-\infty}^{\infty}\mathrm{d}t' f_j(t')\left[1-\mathrm{erf}\left(\frac{t'}{\sqrt{2}\sigma}\right)\right]\int_{-\infty}^{\infty}\mathrm{d}u f_j(u)\left[1-\mathrm{erf}\left(\frac{u}{\sqrt{2}\sigma}\right)\right]\\
=&\frac{A^2}{4}\left\{\int_{-\infty}^{\infty}\mathrm{d}t' f_j(t')\left[1-\mathrm{erf}\left(\frac{t'}{\sqrt{2}\sigma}\right)\right]\right\}^2\\
=&\frac{A^2}{4}\left\{\int_{-\infty}^{\infty}\mathrm{d}t' f_j(t')-\int_{-\infty}^{\infty}\mathrm{d}t' f_j(t')\mathrm{erf}\left(\frac{t'}{\sqrt{2}\sigma}\right)\right\}^2. \qquad (8)
\end{aligned}
$$

The first integral is $\int_{-\infty}^{\infty}\mathrm{d}t' f_j(t') = A$. The second integral can be solved by taking the derivative with respect to $T_{ij}$:

$$
\begin{aligned}
&\int_{-\infty}^{\infty}\mathrm{d}t' f_j(t')\,\mathrm{erf}\left(\frac{t'}{\sqrt{2}\sigma}\right)\\
=&\int \mathrm{d}T_{ij}\int_{-\infty}^{\infty}\mathrm{d}t'\frac{\mathrm{d}}{\mathrm{d}T_{ij}}f_j(t')\,\mathrm{erf}\left(\frac{t'}{\sqrt{2}\sigma}\right)\\
=&\int \mathrm{d}T_{ij}\int_{-\infty}^{\infty}\mathrm{d}t'\left[-f_j'(t')\right]\,\mathrm{erf}\left(\frac{t'}{\sqrt{2}\sigma}\right)\\
&\text{(because }\frac{\mathrm{d}}{\mathrm{d}T_{ij}}f_j(t') = -\frac{\mathrm{d}}{\mathrm{d}t'}f_j(t') \equiv -f_j'(t')\text{)} \qquad (9)\\
=&\int \mathrm{d}T_{ij}\left\{\left[\mathrm{erf}\left(\frac{t'}{\sqrt{2}\sigma}\right)\left[-f_j(t')\right]\right]_{-\infty}^{\infty}-\int_{-\infty}^{\infty}\mathrm{d}t'\frac{2}{A}f_i(t')\left[-f_j(t')\right]\right\}\\
&\text{(integration by parts)}\\
=&\int \mathrm{d}T_{ij}\left\{\left[1\cdot 0-(-1)\cdot 0\right]+\frac{2}{A}\cdot\frac{A^2}{2\sqrt{\pi}\sigma}\exp\left(-\frac{T_{ij}^2}{4\sigma^2}\right)\right\}\\
=&\frac{A}{\sqrt{\pi}\sigma}\int \mathrm{d}T_{ij}\exp\left(-\frac{T_{ij}^2}{4\sigma^2}\right)\\
=&A\,\mathrm{erf}\left(\frac{T_{ij}}{2\sigma}\right) \qquad (10)
\end{aligned}
$$

Term (1.i) (Equation 8) thus reads:

$$\begin{aligned}&\int_{-\infty}^{\infty}\mathrm{d}t'\int_{-\infty}^{\infty}\mathrm{d}u\int_{0}^{\infty}\mathrm{d}s\int_{0}^{\infty}\mathrm{d}v f_i(t'+s)f_i(u+v)f_j(t')f_j(u)\\ =&\frac{A^2}{4}\left[A-A\,\mathrm{erf}\left(\frac{T_{ij}}{2\sigma}\right)\right]^2=\frac{A^4}{4}\left[1-\mathrm{erf}\left(\frac{T_{ij}}{2\sigma}\right)\right]^2\end{aligned}$$

For (2.i) we find

$$\begin{aligned}&\mp\int_{-\infty}^{\infty}\mathrm{d}t'\int_{-\infty}^{\infty}\mathrm{d}u\int_{0}^{\infty}\mathrm{d}s\int_{-\infty}^{0}\mathrm{d}v f_i(t'+s)f_i(u+v)f_j(t')f_j(u)\\ =&\mp\int_{-\infty}^{\infty}\mathrm{d}t' f_j(t')\int_{-\infty}^{\infty}\mathrm{d}u f_j(u)\int_{0}^{\infty}\mathrm{d}s f_i(t'+s)\int_{-\infty}^{0}\mathrm{d}v f_i(u+v)\\ =&\mp\frac{A^2}{4}\int_{-\infty}^{\infty}\mathrm{d}t' f_j(t')\left[1-\mathrm{erf}\left(\frac{t'}{\sqrt{2}\sigma}\right)\right]\int_{-\infty}^{\infty}\mathrm{d}u f_j(u)\left[1+\mathrm{erf}\left(\frac{u}{\sqrt{2}\sigma}\right)\right]\\ \overset{(10)}{=}&\mp\frac{A^2}{4}\left[A-A\,\mathrm{erf}\left(\frac{T_{ij}}{2\sigma}\right)\right]\left[A+A\,\mathrm{erf}\left(\frac{T_{ij}}{2\sigma}\right)\right]\\ =&\mp\frac{A^4}{4}\left[1-\mathrm{erf}^2\left(\frac{T_{ij}}{2\sigma}\right)\right]\end{aligned}$$

Term (3.i) is symmetric to (2.i) and thus yields the same result. For (4.i) we find (in analogy to the term (1.i)):

$$\begin{aligned}&\int_{-\infty}^{\infty}\mathrm{d}t'\int_{-\infty}^{\infty}\mathrm{d}u\int_{-\infty}^{0}\mathrm{d}s\int_{-\infty}^{0}\mathrm{d}v f_i(t'+s)f_i(u+v)f_j(t')f_j(u)\\ =&\frac{A^2}{4}\left(\int_{-\infty}^{\infty}\mathrm{d}t' f_j(t')\left[1+\mathrm{erf}\left(\frac{t'}{\sqrt{2}\sigma}\right)\right]\right)^2\\ =&\frac{A^4}{4}\left[1+\mathrm{erf}\left(\frac{T_{ij}}{2\sigma}\right)\right]^2\end{aligned}$$

We sum the contributions (1.i) to (4.i) for the odd learning window:

$$\begin{aligned}&\frac{A^4}{4}\left[1-\mathrm{erf}\left(\frac{T_{ij}}{2\sigma}\right)\right]^2-2\cdot\frac{A^4}{4}\left[1-\mathrm{erf}^2\left(\frac{T_{ij}}{2\sigma}\right)\right]+\frac{A^4}{4}\left[1+\mathrm{erf}\left(\frac{T_{ij}}{2\sigma}\right)\right]^2\\ =&\frac{A^4}{4}\left[1-2\,\mathrm{erf}\left(\frac{T_{ij}}{2\sigma}\right)+\mathrm{erf}^2\left(\frac{T_{ij}}{2\sigma}\right)-2+2\,\mathrm{erf}^2\left(\frac{T_{ij}}{2\sigma}\right)+1+2\,\mathrm{erf}\left(\frac{T_{ij}}{2\sigma}\right)+\mathrm{erf}^2\left(\frac{T_{ij}}{2\sigma}\right)\right]\\ =&A^4\,\mathrm{erf}^2\left(\frac{T_{ij}}{2\sigma}\right)\end{aligned} \tag{11}$$

Let us continue with the second term of $F$, which is labeled by '(ii)', and consider the first (of four) integrals in Equation 7, i.e. we continue with contribution (1.ii):

$$
\begin{aligned}
&\int_{-\infty}^{\infty} \mathrm{d}t' \int_{-\infty}^{\infty} \mathrm{d}u \int_{0}^{\infty} \mathrm{d}s \int_{0}^{\infty} \mathrm{d}v f_i(t'+s) f_i(u+v) f_j(t') \delta(t'-u) \\
=&\int_{-\infty}^{\infty} \mathrm{d}t' f_j(t') \int_{-\infty}^{\infty} \mathrm{d}u\, \delta(t'-u) \int_{0}^{\infty} \mathrm{d}s f_i(t'+s) \int_{0}^{\infty} \mathrm{d}v f_i(u+v) \\
=&\frac{A^2}{4} \int_{-\infty}^{\infty} \mathrm{d}t' f_j(t') \left[1-\mathrm{erf}\left(\frac{t'}{\sqrt{2}\sigma}\right)\right] \int_{-\infty}^{\infty} \mathrm{d}u\, \delta(t'-u) \left[1-\mathrm{erf}\left(\frac{u}{\sqrt{2}\sigma}\right)\right] \\
=&\frac{A^2}{4} \int_{-\infty}^{\infty} \mathrm{d}t' f_j(t') \left[1-2\,\mathrm{erf}\left(\frac{t'}{\sqrt{2}\sigma}\right)+\mathrm{erf}^2\left(\frac{t'}{\sqrt{2}\sigma}\right)\right] \\
\overset{(10)}{=}&\frac{A^3}{4}\left[1-2\,\mathrm{erf}\left(\frac{T_{ij}}{2\sigma}\right)+\frac{C}{A}\right],
\end{aligned}
$$

with

$$
C = \int_{-\infty}^{\infty} \mathrm{d}t' f_j(t')\, \mathrm{erf}^2\left(\frac{t'}{\sqrt{2}\sigma}\right),
$$

which will be solved later for special cases. Note that $C$ depends on $T_{ij}$ because $f_j(t')$ depends on $T_{ij}$. For (2.ii) we find:

$$
\begin{aligned}
&\mp \int_{-\infty}^{\infty} \mathrm{d}t' \int_{-\infty}^{\infty} \mathrm{d}u \int_{0}^{\infty} \mathrm{d}s \int_{-\infty}^{0} \mathrm{d}v f_i(t'+s) f_i(u+v) f_j(t') \delta(t'-u) \\
=&\mp \frac{A^2}{4} \int_{-\infty}^{\infty} \mathrm{d}t' f_j(t') \left[1-\mathrm{erf}\left(\frac{t'}{\sqrt{2}\sigma}\right)\right] \left[1+\mathrm{erf}\left(\frac{t'}{\sqrt{2}\sigma}\right)\right] \\
=&\mp \frac{A^2}{4} \int_{-\infty}^{\infty} \mathrm{d}t' f_j(t') \left[1-\mathrm{erf}^2\left(\frac{t'}{\sqrt{2}\sigma}\right)\right] \\
=&\mp \frac{A^3}{4}\left(1-\frac{C}{A}\right).
\end{aligned}
$$

For (3.ii) we find the same:

$$
\begin{aligned}
&\mp \int_{-\infty}^{\infty} \mathrm{d}t' \int_{-\infty}^{\infty} \mathrm{d}u \int_{-\infty}^{0} \mathrm{d}s \int_{0}^{\infty} \mathrm{d}v f_i(t'+s) f_i(u+v) f_j(t') \delta(t'-u) \\
=&\mp \frac{A^2}{4} \int_{-\infty}^{\infty} \mathrm{d}t' f_j(t') \left[1+\mathrm{erf}\left(\frac{t'}{\sqrt{2}\sigma}\right)\right] \left[1-\mathrm{erf}\left(\frac{t'}{\sqrt{2}\sigma}\right)\right] \\
=&\mp \frac{A^3}{4}\left(1-\frac{C}{A}\right).
\end{aligned}
$$

For (4.ii) we find:

$$
\begin{aligned}
&\int_{-\infty}^{\infty} \mathrm{d}t' \int_{-\infty}^{\infty} \mathrm{d}u \int_{-\infty}^{0} \mathrm{d}s \int_{-\infty}^{0} \mathrm{d}v f_i(t'+s) f_i(u+v) f_j(t') \delta(t'-u) \\
=&\frac{A^2}{4} \int_{-\infty}^{\infty} \mathrm{d}t' f_j(t') \left[1 + \operatorname{erf}\left(\frac{t'}{\sqrt{2}\sigma}\right)\right]^2 \\
=&\frac{A^2}{4} \int_{-\infty}^{\infty} \mathrm{d}t' f_j(t') \left[1 + 2\operatorname{erf}\left(\frac{t'}{\sqrt{2}\sigma}\right) + \operatorname{erf}^2\left(\frac{t'}{\sqrt{2}\sigma}\right)\right] \\
=&\frac{A^3}{4} \left[1 + 2\operatorname{erf}\left(\frac{T_{ij}}{2\sigma}\right) + \frac{C}{A}\right].
\end{aligned}
$$

Summing contributions (1.ii) to (4.ii) for the odd window yields:

$$
\begin{aligned}
&\frac{A^3}{4} \left[1 - 2\operatorname{erf}\left(\frac{T_{ij}}{2\sigma}\right) + \frac{C}{A}\right] - 2 \cdot \frac{A^3}{4} \left(1 - \frac{C}{A}\right) + \frac{A^3}{4} \left[1 + 2\operatorname{erf}\left(\frac{T_{ij}}{2\sigma}\right) + \frac{C}{A}\right] \\
=&\frac{A^3}{4} \cdot \frac{4\,C}{A} = CA^2
\end{aligned}
\tag{12}
$$

We continue with contribution (1.iii):

$$
\begin{aligned}
&\int_{-\infty}^{\infty} \mathrm{d}t' \int_{-\infty}^{\infty} \mathrm{d}u \int_{0}^{\infty} \mathrm{d}s \int_{0}^{\infty} \mathrm{d}v f_j(t') f_j(u) f_i(t'+s) \delta(t'+s-u-v) \\
=&\int_{-\infty}^{\infty} \mathrm{d}t' f_j(t') \int_{-\infty}^{\infty} \mathrm{d}u f_j(u) \int_{0}^{\infty} \mathrm{d}s f_i(t'+s) \int_{0}^{\infty} \mathrm{d}v\, \delta(t'+s-u-v)
\end{aligned}
$$

Contribution (1.iii) is non-zero if the argument $t'+s-u-v$ of the delta function in the last integral (across $v$) is zero for some $v$, which varies from 0 to $\infty$. The argument of the delta function is thus zero for some $v$ if $0 \leq t'+s-u < \infty$, which we can rewrite as $u \leq t'+s$ and then use it in the integral across $u$, which leads to

$$\begin{aligned}
&\int_{-\infty}^{\infty} \mathrm{d}t' f_j(t') \int_0^{\infty} \mathrm{d}s f_i(t'+s) \int_{-\infty}^{t'+s} \mathrm{d}u\, f_j(u) \\
=&\frac{A}{2} \int_{-\infty}^{\infty} \mathrm{d}t' f_j(t') \int_0^{\infty} \mathrm{d}s f_i(t'+s) \left[1 + \operatorname{erf}\left(\frac{s+t'-T_{ij}}{\sqrt{2}\sigma}\right)\right] \\
=&\frac{A}{2} \int_{-\infty}^{\infty} \mathrm{d}t' f_j(t') \left[\int_0^{\infty} \mathrm{d}s f_i(t'+s) + \int_0^{\infty} \mathrm{d}s f_i(t'+s) \operatorname{erf}\left(\frac{s+t'-T_{ij}}{\sqrt{2}\sigma}\right)\right] \\
=&\frac{A}{2} \int_{-\infty}^{\infty} \mathrm{d}t' f_j(t') \left[\frac{A}{2}\left(1 - \operatorname{erf}\left(\frac{t'}{\sqrt{2}\sigma}\right)\right) + \int_0^{\infty} \mathrm{d}s f_i(t'+s) \operatorname{erf}\left(\frac{s+t'-T_{ij}}{\sqrt{2}\sigma}\right)\right] \\
=&\frac{A^2}{4} \int_{-\infty}^{\infty} \mathrm{d}t' f_j(t') \left[1 - \operatorname{erf}\left(\frac{t'}{\sqrt{2}\sigma}\right)\right] + \frac{A}{2} \int_{-\infty}^{\infty} \mathrm{d}t' f_j(t') \int_0^{\infty} \mathrm{d}s f_i(t'+s) \operatorname{erf}\left(\frac{s+t'-T_{ij}}{\sqrt{2}\sigma}\right) \\
\overset{(10)}{=}&\frac{A^3}{4} \left[1 - \operatorname{erf}\left(\frac{T_{ij}}{2\sigma}\right)\right] + \frac{D \cdot A}{2}
\end{aligned}$$

with $D := \int_{-\infty}^{\infty} \mathrm{d}t' f_j(t') \int_0^{\infty} \mathrm{d}s f_i(t'+s) \operatorname{erf}\left(\frac{s+t'-T_{ij}}{\sqrt{2}\sigma}\right)$. $D$ will be evaluated later for special cases.
Similarly to (1.iii), we treat (2.iii):

$$\begin{aligned}
&\mp \int_{-\infty}^{\infty} \mathrm{d}t' \int_{-\infty}^{\infty} \mathrm{d}u \int_0^{\infty} \mathrm{d}s \int_{-\infty}^{0} \mathrm{d}v f_j(t') f_j(u) f_i(t'+s) \delta(t'+s-u-v) \\
=&\mp \int_{-\infty}^{\infty} \mathrm{d}t' f_j(t') \int_0^{\infty} \mathrm{d}s f_i(t'+s) \int_{t'+s}^{\infty} \mathrm{d}u f_j(u) \\
=&\mp \frac{A}{2} \int_{-\infty}^{\infty} \mathrm{d}t' f_j(t') \int_0^{\infty} \mathrm{d}s f_i(t'+s) \left[1 - \operatorname{erf}\left(\frac{s+t'-T_{ij}}{\sqrt{2}\sigma}\right)\right] \\
=&\mp \frac{A}{2} \int_{-\infty}^{\infty} \mathrm{d}t' f_j(t') \int_0^{\infty} \mathrm{d}s f_i(t'+s) \pm \frac{A}{2} \int_{-\infty}^{\infty} \mathrm{d}t' f_j(t') \int_0^{\infty} \mathrm{d}s f_i(t'+s) \operatorname{erf}\left(\frac{s+t'-T_{ij}}{\sqrt{2}\sigma}\right) \\
=&\mp \frac{A^2}{4} \int_{-\infty}^{\infty} \mathrm{d}t' f_j(t') \left[1 - \operatorname{erf}\left(\frac{t'}{\sqrt{2}\sigma}\right)\right] \pm \frac{A}{2} \int_{-\infty}^{\infty} \mathrm{d}t' f_j(t') \int_0^{\infty} \mathrm{d}s f_i(t'+s) \operatorname{erf}\left(\frac{s+t'-T_{ij}}{\sqrt{2}\sigma}\right) \\
=&\mp \frac{A^3}{4} \left[1 - \operatorname{erf}\left(\frac{T_{ij}}{2\sigma}\right)\right] \pm \frac{D \cdot A}{2}
\end{aligned}$$

For (3.iii) we find:

$$
\begin{aligned}
&\mp \int_{-\infty}^{\infty} \mathrm{d}t' \int_{-\infty}^{\infty} \mathrm{d}u \int_{-\infty}^{0} \mathrm{d}s \int_{0}^{\infty} \mathrm{d}v f_j(t') f_j(u) f_i(t'+s) \delta(t'+s-u-v) \\
&= \mp \int_{-\infty}^{\infty} \mathrm{d}t' f_j(t') \int_{-\infty}^{0} \mathrm{d}s f_i(t'+s) \int_{-\infty}^{t'+s} \mathrm{d}u f_j(u) \\
&= \mp \frac{A}{2} \int_{-\infty}^{\infty} \mathrm{d}t' f_j(t') \int_{-\infty}^{0} \mathrm{d}s f_i(t'+s) \left[1 + \mathrm{erf}\left(\frac{s+t'-T_{ij}}{\sqrt{2}\sigma}\right)\right] \\
&= \mp \frac{A^2}{4} \int_{-\infty}^{\infty} \mathrm{d}t' f_j(t') \left[1 + \mathrm{erf}\left(\frac{t'}{\sqrt{2}\sigma}\right)\right] \mp \frac{A}{2} \int_{-\infty}^{\infty} \mathrm{d}t' f_j(t') \int_{-\infty}^{0} \mathrm{d}s f_i(t'+s)\, \mathrm{erf}\left(\frac{s+t'-T_{ij}}{\sqrt{2}\sigma}\right) \\
&= \mp \frac{A^3}{4} \left[1 + \mathrm{erf}\left(\frac{T_{ij}}{2\sigma}\right)\right] \mp \frac{D' \cdot A}{2}\,.
\end{aligned}
$$

with $D' := \int_{-\infty}^{\infty} \mathrm{d}t' f_j(t') \int_{-\infty}^{0} \mathrm{d}s f_i(t'+s)\, \mathrm{erf}\left(\frac{s+t'-T_{ij}}{\sqrt{2}\sigma}\right)$, which we will evaluate later for special cases.

Finally, for (4.iii) we find

$$
\begin{aligned}
&\int_{-\infty}^{\infty} \mathrm{d}t' \int_{-\infty}^{\infty} \mathrm{d}u \int_{-\infty}^{0} \mathrm{d}s \int_{-\infty}^{0} \mathrm{d}v f_j(t') f_j(u) f_i(t'+s) \delta(t'+s-u-v) \\
&= \int_{-\infty}^{\infty} \mathrm{d}t' f_j(t') \int_{-\infty}^{0} \mathrm{d}s f_i(t'+s) \int_{t'+s}^{\infty} \mathrm{d}u f_j(u) \\
&= \frac{A}{2} \int_{-\infty}^{\infty} \mathrm{d}t' f_j(t') \int_{-\infty}^{0} \mathrm{d}s f_i(t'+s) \left[1 - \mathrm{erf}\left(\frac{s+t'-T_{ij}}{\sqrt{2}\sigma}\right)\right] \\
&= \frac{A^2}{4} \int_{-\infty}^{\infty} \mathrm{d}t' f_j(t') \left[1 + \mathrm{erf}\left(\frac{t'}{\sqrt{2}\sigma}\right)\right] - \frac{A}{2} \int_{-\infty}^{\infty} \mathrm{d}t' f_j(t') \int_{-\infty}^{0} \mathrm{d}s f_i(t'+s)\, \mathrm{erf}\left(\frac{s+t'-T_{ij}}{\sqrt{2}\sigma}\right) \\
&= \frac{A^3}{4} \left[1 + \mathrm{erf}\left(\frac{T_{ij}}{2\sigma}\right)\right] - \frac{D' \cdot A}{2}\,.
\end{aligned}
$$

To sum the four contributions (1.iii) to (4.iii) for the odd window, we note that the first terms (square brackets) of (1.iii) and (2.iii) cancel, as well as the first terms of (3.iii) and (4.iii). We thus obtain:

$$
\frac{D \cdot A}{2} + \frac{D \cdot A}{2} - \frac{D' \cdot A}{2} - \frac{D' \cdot A}{2} = A(D - D')\,.
$$

We continue with contribution (1.iv):

$$
\begin{aligned}
&\int_{-\infty}^{\infty} \mathrm{d}t' \int_{-\infty}^{\infty} \mathrm{d}u \int_0^{\infty} \mathrm{d}s \int_0^{\infty} \mathrm{d}v f_i(t'+s)\delta(t'+s-u-v) f_j(t')\delta(t'-u) \\
=&\int_{-\infty}^{\infty} \mathrm{d}t' f_j(t') \int_{-\infty}^{\infty} \mathrm{d}u\, \delta(t'-u) \int_0^{\infty} \mathrm{d}s f_i(t'+s) \int_0^{\infty} \mathrm{d}v\, \delta(t'+s-u-v) \\
=&\int_{-\infty}^{\infty} \mathrm{d}t' f_j(t') \int_0^{\infty} \mathrm{d}s f_i(t'+s) \int_{-\infty}^{t'+s} \mathrm{d}u\, \delta(t'-u) \\
\overset{u'=u-t'}{=}&\int_{-\infty}^{\infty} \mathrm{d}t' f_j(t') \int_0^{\infty} \mathrm{d}s f_i(t'+s) \int_{-\infty}^{s} \mathrm{d}u'\, \delta(u') \\
=&\int_{-\infty}^{\infty} \mathrm{d}t' f_j(t') \int_0^{\infty} \mathrm{d}s f_i(t'+s) \cdot \begin{cases} 1, \text{for } s>0 \\ 0, \text{else} \end{cases} \\
=&\int_{-\infty}^{\infty} \mathrm{d}t' f_j(t') \int_0^{\infty} \mathrm{d}s f_i(t'+s) \\
=&\frac{A}{2} \int_{-\infty}^{\infty} \mathrm{d}t' f_j(t') \left[1-\mathrm{erf}\left(\frac{t'}{\sqrt{2}\sigma}\right)\right] \\
\overset{(10)}{=}&\frac{A^2}{2}\left[1-\mathrm{erf}\left(\frac{T_{ij}}{2\sigma}\right)\right]
\end{aligned}
$$

By similar arguments, (2.iv) yields:

$$
\begin{aligned}
&\mp \int_{-\infty}^{\infty} \mathrm{d}t' \int_{-\infty}^{\infty} \mathrm{d}u \int_0^{\infty} \mathrm{d}s \int_{-\infty}^{0} \mathrm{d}v f_i(t'+s)\delta(t'+s-u-v) f_j(t')\delta(t'-u) \\
=&\mp \int_{-\infty}^{\infty} \mathrm{d}t' f_j(t') \int_0^{\infty} \mathrm{d}s f_i(t'+s) \int_{t'+s}^{\infty} \mathrm{d}u\, \delta(t'-u) \\
=&\mp \int_{-\infty}^{\infty} \mathrm{d}t' f_j(t') \int_0^{\infty} \mathrm{d}s f_i(t'+s) \int_{s}^{\infty} \mathrm{d}u'\, \delta(u') \\
=&\mp \int_{-\infty}^{\infty} \mathrm{d}t' f_j(t') \int_0^{\infty} \mathrm{d}s f_i(t'+s) \cdot \begin{cases} 1, \text{for } s<0 \\ 0, \text{else} \end{cases} \\
=&0\,.
\end{aligned}
$$

(3.iv) yields

$$
\begin{aligned}
&\mp \int_{-\infty}^{\infty} \mathrm{d}t' \int_{-\infty}^{\infty} \mathrm{d}u \int_{-\infty}^{0} \mathrm{d}s \int_{0}^{\infty} \mathrm{d}v f_i(t'+s)\delta(t'+s-u-v) f_j(t')\delta(t'-u) \\
=&\mp \int_{-\infty}^{\infty} \mathrm{d}t' f_j(t') \int_{-\infty}^{0} \mathrm{d}s f_i(t'+s) \int_{-\infty}^{t'+s} \mathrm{d}u\, \delta(t'-u) \\
=&\mp \int_{-\infty}^{\infty} \mathrm{d}t' f_j(t') \int_{-\infty}^{0} \mathrm{d}s f_i(t'+s) \cdot \begin{cases} 1, \text{for } s>0 \\ 0, \text{else} \end{cases} \\
=&0\,.
\end{aligned}
$$

(4.iv) yields

$$
\begin{aligned}
&\int_{-\infty}^{\infty} \mathrm{d}t' \int_{-\infty}^{\infty} \mathrm{d}u \int_{-\infty}^{0} \mathrm{d}s \int_{-\infty}^{0} \mathrm{d}v f_i(t'+s)\delta(t'+s-u-v) f_j(t')\delta(t'-u) \\
=&\int_{-\infty}^{\infty} \mathrm{d}t' f_j(t') \int_{-\infty}^{0} \mathrm{d}s f_i(t'+s) \int_{t'+s}^{\infty} \mathrm{d}u\, \delta(t'-u) \\
=&\int_{-\infty}^{\infty} \mathrm{d}t' f_j(t') \int_{-\infty}^{0} \mathrm{d}s f_i(t'+s) \cdot \begin{cases} 1, \text{for } s<0 \\ 0, \text{else} \end{cases} \\
=&\int_{-\infty}^{\infty} \mathrm{d}t' f_j(t') \int_{-\infty}^{0} \mathrm{d}s f_i(t'+s) \\
=&\frac{A}{2} \int_{-\infty}^{\infty} \mathrm{d}t' f_j(t') \left[1+\mathrm{erf}\left(\frac{t'}{\sqrt{2}\sigma}\right)\right] \\
\overset{(10)}{=}&\frac{A^2}{2}\left[1+\mathrm{erf}\left(\frac{T_{ij}}{2\sigma}\right)\right]\,.
\end{aligned}
$$

We sum the contributions (1.iv) and (4.iv) and obtain $A^2$.
We now collect all terms for the odd window:

$$
\frac{1}{\mu^2}\langle \Delta w_{ij}^2 \rangle = A^4 \mathrm{erf}^2\left(\frac{T_{ij}}{2\sigma}\right) + CA^2 + A(D-D') + A^2.
$$

So far we have calculated the second moment of $\Delta w_{ij}$. In order to determine the variance, we need to calculate the average weight change for the odd window

$$
\begin{aligned}
\langle \Delta w_{ij} \rangle =& \int_{-\infty}^{\infty} \mathrm{d}t \, W(t) C_{ij}(t) \\
=& \mu \int_{-\infty}^{\infty} \mathrm{d}t \, \mathrm{sgn}(t) C_{ij}(t) \\
=& \mu \int_{-\infty}^{\infty} \mathrm{d}t \, \mathrm{sgn}(t) \int_{-\infty}^{\infty} \mathrm{d}t' f_i(t') f_j(t'+t) \\
=& \frac{A^2 \mu}{2\sqrt{\pi}\sigma} \int_{-\infty}^{\infty} \mathrm{d}t \, \mathrm{sgn}(t) \exp\left(-\frac{(t-T_{ij})^2}{4\sigma^2}\right) \quad \text{(using Eq. 15 of the main text)} \\
=& \frac{A^2 \mu}{2\sqrt{\pi}\sigma} \left[\int_{0}^{\infty} \mathrm{d}t \, \exp\left(-\frac{(t-T_{ij})^2}{4\sigma^2}\right) - \int_{-\infty}^{0} \mathrm{d}t \, \exp\left(-\frac{(t-T_{ij})^2}{4\sigma^2}\right)\right] \\
=& \frac{A^2 \mu}{2} \left[1 + \mathrm{erf}\left(\frac{T_{ij}}{2\sigma}\right) - \left(1 - \mathrm{erf}\left(\frac{T_{ij}}{2\sigma}\right)\right)\right] \\
=& A^2 \mu \, \mathrm{erf}\left(\frac{T_{ij}}{2\sigma}\right) . \qquad (13)
\end{aligned}
$$

The variance thus reads:

$$
\begin{aligned}
\frac{1}{\mu^2} \mathrm{var}(\Delta w_{ij}) =& \langle \Delta w_{ij}^2 \rangle - \langle \Delta w_{ij} \rangle^2 \\
=& A^4 \mathrm{erf}^2\left(\frac{T_{ij}}{2\sigma}\right) + CA^2 + A(D-D') + A^2 - A^4 \mathrm{erf}^2\left(\frac{T_{ij}}{2\sigma}\right) \\
=& (C+1)A^2 + A(D-D') . \qquad (14)
\end{aligned}
$$

For the signal-to-noise ratio we note that the definition from Equation 1, for odd learning windows, simplifies to

$$
\mathrm{SNR} = \frac{\langle \Delta w_{ij} \rangle - \langle \Delta w_{ji} \rangle}{\mathrm{std}\,(\Delta w_{ij}) + \mathrm{std}\,(\Delta w_{ji})} = \frac{\langle \Delta w_{ij} \rangle}{\mathrm{std}\,(\Delta w_{ij})} ,
$$

because $\Delta w_{ij} = -\Delta w_{ji}$ for odd learning windows.

We insert Equations 13 and 14 and find

$$
\mathrm{SNR} = \frac{A^2 \, \mathrm{erf}\left(\frac{T_{ij}}{2\sigma}\right)}{\sqrt{(C+1)A^2 + A(D-D')}} .
$$

To obtain the final result, we have to evaluate $C$, $D$, and $D'$. We distinguish the two cases $T_{ij} \gg \sigma$ and $T_{ij} = \sigma$ to approximate these three terms:

1. $T_{ij} \gg \sigma$:

$$\mathrm{erf}\left(\frac{T_{ij}}{2\sigma}\right) \approx 1 \tag{15}$$

$$C = \int_{-\infty}^{\infty} \mathrm{d}t' f_j(t') \mathrm{erf}^2\left(\frac{t'}{\sqrt{2}\sigma}\right) \approx A\,,$$

because the Gaussian function $f_j$ is shifted far into the positive lobe of the error function.

$$\begin{aligned} D &= \int_{-\infty}^{\infty} \mathrm{d}t' f_j(t') \int_{0}^{\infty} \mathrm{d}s f_i(t'+s)\,\mathrm{erf}\left(\frac{t'+s-T_{ij}}{\sqrt{2}\sigma}\right) \\ &\approx \int_{-\infty}^{\infty} \mathrm{d}t' f_j(t') \int_{0}^{\infty} \mathrm{d}s\,[-f_i(t'+s)] \\ &= -\frac{A}{2}\int_{-\infty}^{\infty} \mathrm{d}t' f_j(t') \left[1 - \mathrm{erf}\left(\frac{t'}{\sqrt{2}\sigma}\right)\right] \\ &\overset{(10)}{=} -\frac{A}{2}\left[A - A\,\mathrm{erf}\left(\frac{T_{ij}}{2\sigma}\right)\right] \\ &\overset{(15)}{\approx} -\frac{A}{2}\,[A-A] \\ &= 0\,. \end{aligned}$$

$$\begin{aligned} D' &= \int_{-\infty}^{\infty} \mathrm{d}t' f_j(t') \int_{-\infty}^{0} \mathrm{d}s f_i(t'+s)\,\mathrm{erf}\left(\frac{t'+s-T_{ij}}{\sqrt{2}\sigma}\right) \\ &= -\frac{A}{2}\int_{-\infty}^{\infty} \mathrm{d}t' f_j(t') \left[1 + \mathrm{erf}\left(\frac{t'}{\sqrt{2}\sigma}\right)\right] \\ &\overset{(10),(15)}{\approx} -\frac{A}{2}\,[A+A] \\ &= -A^2\,. \end{aligned}$$

Thus,

$$\mathrm{SNR} \approx \frac{A^2 \cdot 1}{\sqrt{(A+1)A^2 + A(0+A^2)}} = \frac{A^2}{\sqrt{2A^3+A^2}} = \frac{A}{\sqrt{2A+1}}\,. \tag{16}$$

This number (for $A = 10$) is indicated as the analytical comparison in Figure 5E. For large A, the SNR (expression 16) approaches $\sqrt{A/2}$.

2. $T_{ij} = \sigma$:

$$\mathrm{erf}\left(\frac{T_{ij}}{2\sigma}\right) = \mathrm{erf}\left(\frac{1}{2}\right) \approx 0.52$$

$$C \approx 0.494\,A,$$
$$D \approx -0.013\,A^2\,,$$
$$D' \approx -0.507\,A^2\,,$$

all of which we calculated numerically.

It follows:

$$\begin{aligned} \mathrm{SNR} &\approx \frac{A^2\,\mathrm{erf}\left(\frac{1}{2}\right)}{\sqrt{(C+1)A^2 + A(D-D')}} \\ &\approx \frac{0.52\,A^2}{\sqrt{0.494\,A^3 + A^2 + 0.494\,A^3}} \\ &\approx \frac{0.52\,A^2}{\sqrt{0.99\,A^3 + A^2}} \\ &\overset{A=10}{\approx} 1.58 \end{aligned} \tag{17}$$

This number is plotted as the large-tau approximation in Figure 4C. For large $A$, we find SNR $\propto \sqrt{A}$.

**Even windows**

As argued in the main text, for even windows, the weight change $\Delta w_{ij}$ contains no information about the order of events because $\Delta w_{ij} = \Delta w_{ji}$. This can be seen from Equation 1 of this supplement. The SNR is zero for purely even windows because the signal is zero. Nonetheless, we can calculate the variance of the weight change. To do so, we collect all terms of $\langle \Delta w_{ij}^2 \rangle$ for even windows (indicated by the bottom symbol of all occurences of '$\pm$' and '$\mp$' in the previous section). Again, we assume wide windows ($\kappa \to \infty$). We find

Collecting the terms (1.i) to (4.i) yields

$$\frac{A^4}{4}\left[1-\mathrm{erf}\left(\frac{T_{ij}}{2\sigma}\right)\right]^2 + 2\cdot\frac{A^4}{4}\left[1-\mathrm{erf}^2\left(\frac{T_{ij}}{2\sigma}\right)\right] + \frac{A^4}{4}\left[1+\mathrm{erf}\left(\frac{T_{ij}}{2\sigma}\right)\right]^2 = A^4\,.$$

Similarly, we sum the terms (1.ii) to (4.ii):

$$\frac{A^3}{4}\left[1-2\,\mathrm{erf}\left(\frac{T_{ij}}{2\sigma}\right)+\frac{C}{A}\right] + 2\cdot\frac{A^3}{4}\left(1-\frac{C}{A}\right) + \frac{A^3}{4}\left[1+2\,\mathrm{erf}\left(\frac{T_{ij}}{2\sigma}\right)+\frac{C}{A}\right] = A^3\,.$$

We continue to collect the contributions (1.iii) to (4.iii):

$$\begin{aligned}
&\frac{A^3}{4}\left[1-\operatorname{erf}\left(\frac{T_{ij}}{2\sigma}\right)\right]+\frac{D\cdot A}{2}\\
+&\frac{A^3}{4}\left[1-\operatorname{erf}\left(\frac{T_{ij}}{2\sigma}\right)\right]-\frac{D\cdot A}{2}\\
+&\frac{A^3}{4}\left[1+\operatorname{erf}\left(\frac{T_{ij}}{2\sigma}\right)\right]+\frac{D'\cdot A}{2}\\
+&\frac{A^3}{4}\left[1+\operatorname{erf}\left(\frac{T_{ij}}{2\sigma}\right)\right]-\frac{D'\cdot A}{2}\\
=&A^3
\end{aligned}$$

Finally, summing (1.iv) to (4.iv) yield the same result as for the odd window: $A^2$.

Taken together,

$$\frac{1}{\lambda^2}\langle\Delta w_{ij}^2\rangle = \frac{1}{\lambda^2}\langle\Delta w_{ji}^2\rangle = A^4 + 2\,A^3 + A^2\,.$$

Together with

$$\frac{1}{\lambda}\langle\Delta w_{ij}\rangle = \frac{1}{\lambda}\langle\Delta w_{ji}\rangle = A^2\,,$$

the variance reads:

$$\frac{1}{\lambda^2}\,\mathrm{var}(\Delta w_{ij}) = \frac{1}{\lambda^2}\,\mathrm{var}(\Delta w_{ji}) = 2\,A^3 + A^2\,.$$

We now insert these variances in the denominator of Equation 1:

$$\frac{1}{\lambda}\left[\mathrm{std}\left(\Delta w_{ij}\right)+\mathrm{std}\left(\Delta w_{ji}\right)\right] = 2\sqrt{2A^3+A^2}\,,$$

which, assuming $\mu = \lambda$, is twice the noise as for odd windows ($\sqrt{2A^3+A^2}$, Equation 16).

In summary, for a complex learning window with even and odd contributions, the signal solely depends on the odd part, whereas both parts, even and odd, contribute to the noise. Any even contribution thus only decreases the SNR.
